# Selective suppression of oligodendrocyte-derived amyloid beta rescues neuronal dysfunction in Alzheimer’s Disease

**DOI:** 10.1101/2024.06.21.600003

**Authors:** Rikesh M. Rajani, Robert Ellingford, Mariam Hellmuth, Samuel S. Harris, Orjona S. Taso, David Graykowski, Francesca Kar Wey Lam, Charles Arber, Emre Fertan, John S. H. Danial, Matthew Swire, Marcus Lloyd, Tatiana A. Giovannucci, Mathieu Bourdenx, David Klenerman, Robert Vassar, Selina Wray, Carlo Sala Frigerio, Marc Aurel Busche

## Abstract

Reduction of amyloid beta (Aβ) has been shown to be effective in treating Alzheimer’s Disease (AD), but the underlying assumption that neurons are the main source of pathogenic Aβ is untested. Here we challenge this prevailing belief by demonstrating that oligodendrocytes are an important source of Aβ, and play a key role in promoting abnormal neuronal hyperactivity in AD. We show that selectively suppressing oligodendrocyte Aβ production improves AD brain pathology and restores neuronal function *in vivo*. Our findings suggest that targeting oligodendrocyte Aβ production could be a promising therapeutic strategy for treating AD.

---

Alzheimer’s Disease (AD) is a devastating neurodegenerative disorder affecting millions of people worldwide. Accumulation of amyloid beta (Aβ) is an early critical hallmark of the disease, and thus is an important target for understanding pathophysiology and therapy. Recent clinical trials demonstrating a slowing of cognitive and functional decline in individuals with AD treated with anti-Aβ antibodies indeed reinforce the important role of Aβ in AD pathophysiology^1,2^. Among the earliest responses of neurons to this accumulation of Aβ is an abnormal increase in excitability^3,4^. However, neurons are not the only cells to react to Aβ. Recently, transcriptomic studies have shown changes not only in microglia and astrocytes but also in oligodendrocytes, the myelinating cells of the central nervous system, in both human AD tissue^5,6^ and in mouse models of AD^7,8^. In addition, genetic risk associated with AD is enriched in age-dependent transcriptome networks of both microglia and oligodendrocytes^9^.

Despite these key cellular effects of Aβ, and its essential role in AD, the traditional assumption that neurons are the primary source of pathogenic Aβ in the brain has remained untested. In this study, we show that oligodendrocytes in human tissue contain all of the components required to produce Aβ, and that human oligodendrocytes produce soluble Aβ *in vitro*. We further show that selectively suppressing oligodendrocyte Aβ production in an AD mouse model is sufficient to rescue abnormal neuronal hyperactivity. Thus, we provide evidence for a critical role of oligodendrocyte-derived Aβ for early neuronal dysfunction in AD.

## Results

### Oligodendrocytes contain all components required to produce Aβ

To identify the cell types in the brain that are intrinsically capable of producing Aβ, we exploited four publicly available human single nucleus RNA sequencing (snRNA-seq) datasets^5,10–12^ and examined expression levels of genes involved in the production of Aβ: the amyloid precursor protein (*APP*), beta secretase (*BACE1*), and components of gamma secretase [presenilin (*PSEN*) 1, *PSEN2*, nicastrin (*NCSTN*), *APH1A*, *APH1B, PSENEN*]. We found that, apart from *PSEN2* (which is not essential for gamma secretase function when *PSEN1* is present^13^), oligodendrocytes had elevated expression of all of these genes (**Fig. 1a-d**). Remarkably, many genes were expressed at a higher level in oligodendrocytes than in any other cell type in the brain, including notably neurons (**Fig. 1a-d**). With the exception of neurons and oligodendrocytes, no other cell type examined was found to highly express all of the genes required for Aβ production (**Fig. 1a-d**), suggesting that oligodendrocytes, uniquely among glia, have an intrinsic capacity to produce Aβ. To confirm that this is not limited to RNA, we exploited publicly available proteomics data derived from isolated mouse brain cells^14^ and found that, at the protein level, oligodendrocytes indeed contain high levels of APP, BACE1 and PSEN1 (**Supplementary Fig. 1**). We further confirmed the presence of APP and BACE1 protein in mouse oligodendrocytes by immunohistochemistry (**Fig. 1e-f; Supplementary Fig. 2**).

**Fig. 1.**
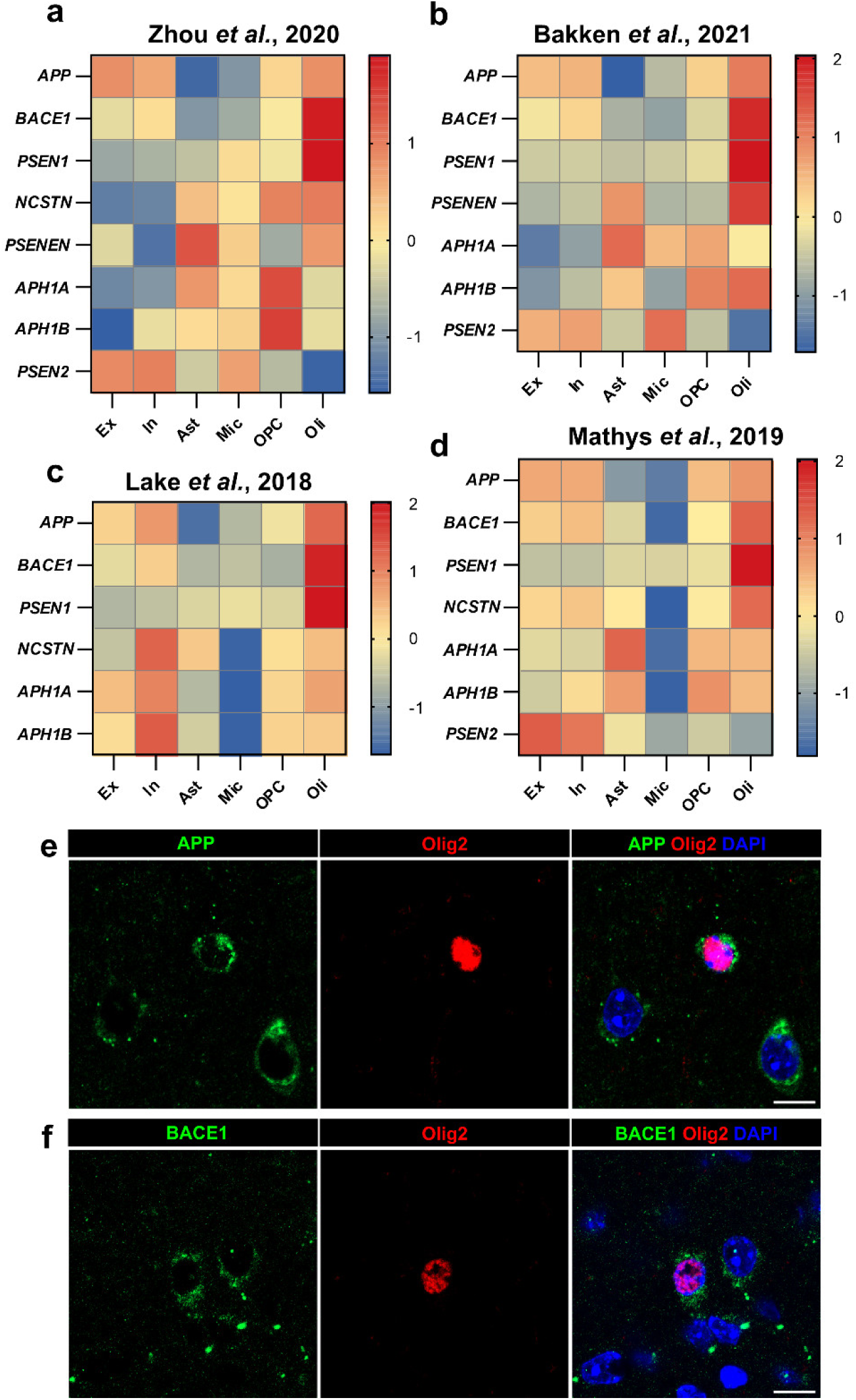
Components required to produce Aβ are expressed at high levels in oligodendrocytes, but not other glial cells. Heatmaps showing the log_2_ (norm count) z-score of genes of interest across different cell types [Excitatory neurons (Ex), Inhibitory neurons (In), Astrocytes (Ast), Microglia (Mic), Oligodendrocyte Precursor Cells (OPC), and Oligodendrocytes (Oli)], from 4 publicly available human single nucleus RNA sequencing datasets. *APP*, *BACE1*, and all components of γ-secretase (*PSEN1*, *PSENEN*, *NCSTN*, *APH1A*, *APH1B*) with the exception of *PSEN2* (which is interchangeable with *PSEN1*) are expressed at high levels in oligodendrocytes, many at higher levels than any other cell type. **a,** Data from Zhou *et al.*, 2020^10^ was generated using tissue from the motor cortex of 36 subjects including controls, AD patients, and those carrying TREM2 variants. **b,** Data from Bakken *et al.*, 2021^11^ was generated using tissue from the motor cortex of 5 control subjects. **c,** Data from Lake *et al.*, 2018^12^ was generated using tissue from the frontal cortex of 6 control subjects. **d,** Data from Mathys *et al.,* 2019^5^ was generated using tissue from the prefrontal cortex of 48 subjects with varying degrees of AD-related pathology. **e,** Representative immunofluorescent images showing APP (green), oligodendroglial marker Olig2 (red) and DAPI (nuclei; blue) in the cortex of a 4-month-old wild type mouse. Scale bar = 10μm. **f,** Representative immunofluorescent images showing BACE1 (green), oligodendroglial marker Olig2 (red) and DAPI (nuclei; blue) in the cortex of a 4-month-old wild type mouse. Scale bar = 10μm.

### Human AD brains possess more oligodendrocytes with the capacity to produce Aβ

To validate these findings, and to determine whether the capacity of oligodendrocytes to produce Aβ is altered in AD, we performed RNAscope *in situ* hybridization (ISH) on post-mortem tissue from the brains of patients with sporadic AD (sAD) and controls for *MBP* (a gene expressed exclusively in oligodendrocytes in the central nervous system^15^), *APP*, and *BACE1* (**Fig. 2a**). We found that ∼80% of oligodendrocytes express both *APP* and *BACE1* in Layers 5/6 of both sAD patient and control brains, indicating that they are capable of producing Aβ (**Fig. 2b**). Contrary to our expectation of oligodendrocyte loss associated with myelin loss in AD^16^, we observed an increased number of oligodendrocytes in Layers 5/6 of the prefrontal cortex of brains from sAD patients (**Fig. 2c**), which was not solely explained by age (**Supplementary Fig. 3a**) and was not seen in Layers 2/3 of the prefrontal cortex (**Supplementary Fig. 3b**). Combined with the equivalent proportion of oligodendrocytes expressing both *APP* and *BACE1*, this increase in oligodendrocyte density resulted in an increased number of Aβ-capable oligodendrocytes within the brains of sAD patients (**Fig. 2d**). By examining all cells expressing both *APP* and *BACE1*, we observed an increase in the total number of *APP*^+^ *BACE1*^+^ cells in the brains of sAD patients (**Supplementary Fig. 3c**), which was driven by an elevated proportion of *APP*^+^ *BACE1*^+^ cells that are oligodendrocytes (**Supplementary Fig. 3d**). Notably, there was no change in the number of other cells (*MBP*^−^) capable of producing Aβ (**Supplementary Fig. 3e**), which were almost exclusively neurons (labelled as *RBFOX3*^+^, a gene expressed in all neurons in the cortex; **Supplementary Fig. 3f-i**).

**Fig. 2:**
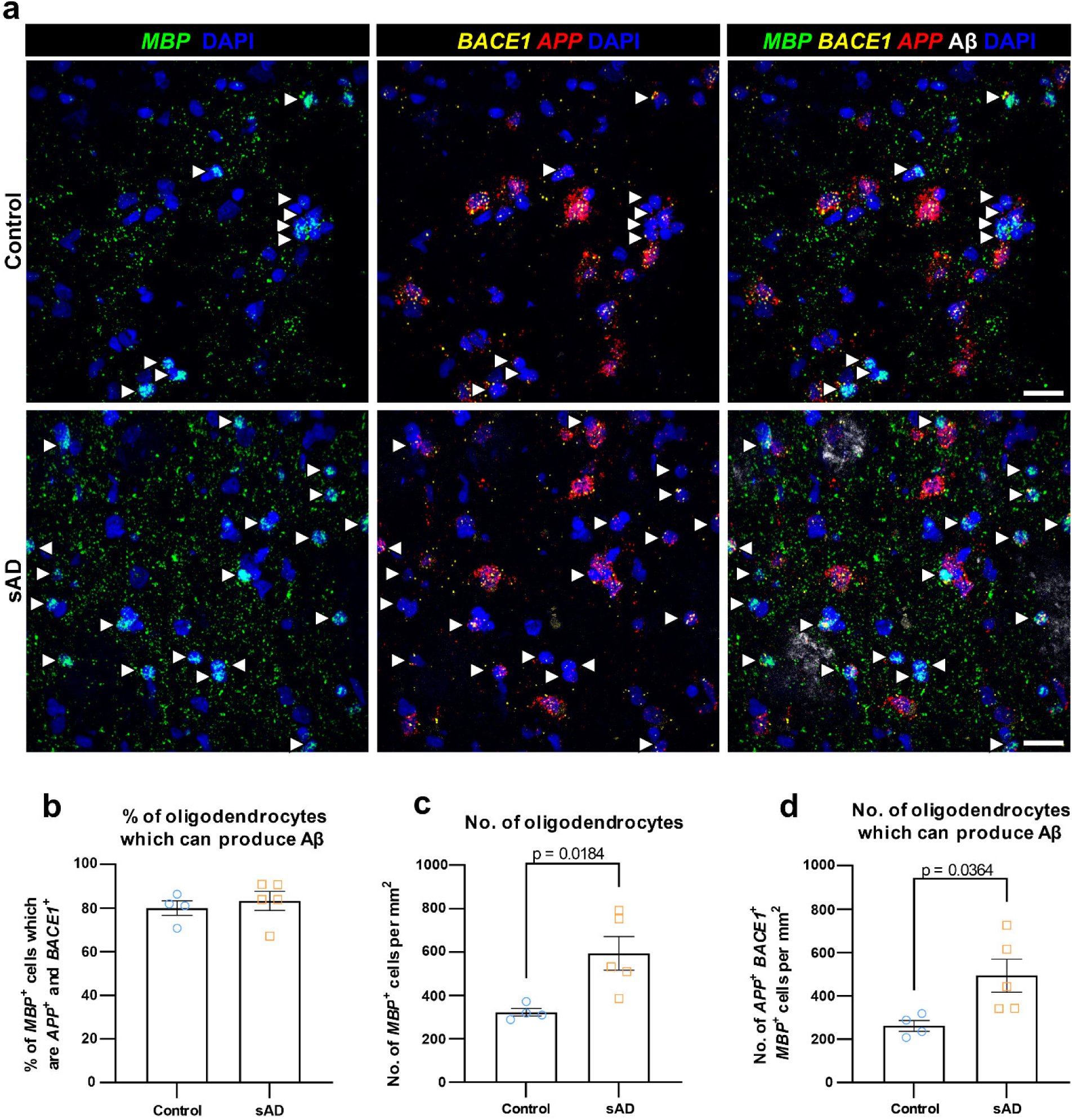
Human sporadic AD brains have more oligodendrocytes capable of producing Aβ compared to controls. **a**, Fluorescence images from Layers 5/6 of control (top) and sporadic AD (sAD; bottom) post-mortem human prefrontal cortex labelled for *MBP* (oligodendrocyte-specific gene; green), *BACE1* (yellow), *APP* (red), Aβ (identified by 6E10-antibody; white), and DAPI (nuclei; blue). Aβ-capable oligodendrocytes (*MBP*^+^ *BACE1*^+^ *APP*^+^ nuclei) are marked with white arrowheads. Scale bar = 25μm **b,** Quantification showing that ∼80% of oligodendrocytes are capable of producing Aβ in both control and sAD brains. **c,** Quantification showing an increase in the number of oligodendrocytes in Layers 5/6 of sAD brains. **d,** Quantification showing significantly more Aβ-capable oligodendrocytes in sAD brains than controls. In **b**,**c,d**, each data point represents a single brain (n=4 control brains, n=5 sAD brains) with bars representing mean ± SEM; unpaired t-test: *t*(7)=0.568, 3.058, 2.581 in **b,c,d** respectively.

We additionally carried out densitometric analysis of *APP* and *BACE1* spots within oligodendrocytes and neurons. On average, oligodendrocytes were found to express higher amounts of *APP* and *BACE1* than neurons (**Supplementary Fig. 4a-b**), though neurons showed a greater variability in expression levels, with a subset of neurons showing very high expression yet others showing only minimal expression (**Supplementary Fig. 4c-d**). The observed increase in the number of Aβ-capable oligodendrocytes in sAD brains suggests that oligodendrocyte Aβ production may play an important, but hitherto unappreciated, role in the pathogenesis of the disease.

### Human iPSC-derived oligodendrocytes produce Aβ

As well as expressing the components necessary for Aβ production, it is pathologically relevant to understand whether human oligodendrocytes do indeed produce Aβ. To determine this, we differentiated oligodendrocytes from human induced pluripotent stem cells (iPSCs) following a previously established protocol^17^. We used iPSCs derived from familial AD (fAD) patients as well as an isogenic control, to examine this question in well-established lines both with and without pathological mutations affecting Aβ production^18,19^; due to the rarity of iPSC lines with fAD mutations we did not compare between different lines, but considered them together. After 30 days of maturation, these cultures contained 6.3% MBP^+^ oligodendrocytes and 87.3% oligodendrocyte precursor cells (OPCs)/pre-myelinating oligodendrocytes, but almost no other non-oligodendroglial cell types (**Fig. 3a; Supplementary Fig. 5a-b**), consistent with previous studies^17^. Across all lines tested we found that human iPSC-derived oligodendrocytes indeed produced Aβ, and that this Aβ production was significantly reduced by administration of the BACE1 inhibitor NB-360^20^, indicating that Aβ production in oligodendrocytes is BACE1 dependent (**Fig. 3b**). Notably, this reduction in Aβ production is similar to what has previously been shown in human iPSC-derived neurons upon administration of NB-360 or similar BACE inhibitors^21–24^; these data from human iPSCs are also consistent with, and translationally extend, previous findings from rat oligodendrocyte cultures^25^.

**Fig. 3:**
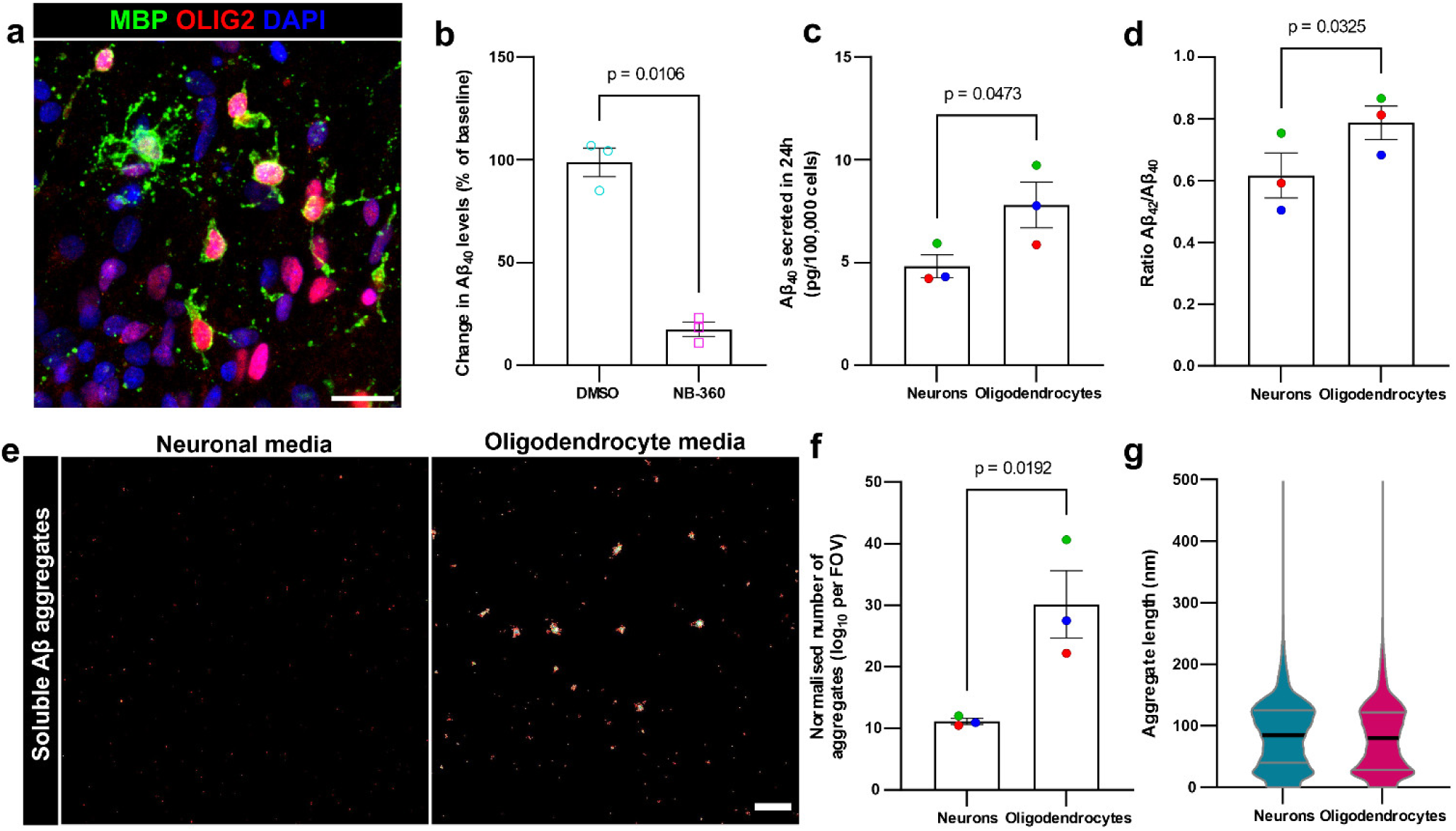
Human oligodendrocytes produce soluble Aβ and aggregates at higher levels than neurons. **a**, Fluorescent image of human iPSC-derived oligodendrocyte culture immunolabelled for MBP (myelin basic protein; green), OLIG2 (marker of all oligodendroglia; red), and DAPI (nuclei; blue). This example shows multiple mature oligodendrocytes extending MBP^+^ myelin processes, while the majority of other cells are OLIG2^+^ MBP^-^ oligodendrocyte precursor cells (OPCs). Scale bar = 25μm. **b,** Quantification by ELISA showing a significant reduction in the amount of Aβ_40_ produced (as a % of the amount produced prior to treatment) by human oligodendrocytes when treated with BACE1 inhibitor (NB-360) compared to vehicle control (DMSO). **c,** ELISA data showing more Aβ_40_ produced by oligodendrocytes than neurons derived from the same familial AD (fAD) human-iPSC lines. **d,** Quantification by ELISA showing higher Aβ_42_/Aβ_40_ ratio produced by oligodendrocytes compared to neurons derived from the same fAD human-iPSC lines. **e,** DNA-PAINT super-resolution images showing more Aβ aggregates in media from oligodendrocytes (right) compared to neurons (left). Scale bar = 1µm. **f,** Quantification showing oligodendrocytes produce a higher proportion of Aβ as aggregates compared to neurons derived from the same fAD human-iPSC lines. **g,** Violin plots showing the length of aggregates produced by neurons and oligodendrocytes (mean ± standard deviation: Neurons 85.1 ± 55.7, Oligodendrocytes 80.7 ± 58.3; n = 18,550 aggregates from 3 neuron lines [three independent inductions per line] and 24,719 aggregates from 3 oligodendrocyte lines [3 independent inductions per line]). In **b**, each data point represents the average of two independent inductions from each of a different cell line (n=3 cell lines), with bars showing mean ± SEM; paired t test: *t*(2)=9.613. In **c,d,f**, each data point represents the average of four (for PSEN1 WT line neurons in **d**) or three independent inductions from each of a different cell line (n=3 cell lines), showing mean ± SEM with each cell line shown in a different colour to highlight pairing (blue: PSEN1 WT; green: PSEN1 int4del; red: PSEN1 R278I); paired t-test: *t*(2)=4.435, 5.411 in **c,d** respectively; ratio paired t-test: *t*(2)=7.117 in **f**.

Next, to understand how oligodendrocyte Aβ production compares to neuronal Aβ production, we generated human iPSC-derived cortical neurons from the same cell lines following a previously established protocol^26^ (**Supplementary Fig. 5c-f**), and normalised the amount of Aβ produced to cell number. Under the experimental conditions tested, we found that oligodendrocytes produce more Aβ than neurons derived from the same cell line, both in lines from fAD patients with PSEN1 mutations and an isogenic control (**Fig. 3c**). We additionally quantified Aβ production by human iPSC-derived OPCs, microglia and astrocytes (**Supplementary Fig. 6a-f**) and found that these cells produce only low amounts of Aβ (**Supplementary Fig. 6g**), consistent with our analyses of snRNA-seq data (**Fig. 1a-d**).

We next compared the Aβ species produced between the cell types, and found that oligodendrocytes had an increased Aβ 42:40 ratio relative to neurons from the same line (**Fig. 3d**). This observation suggests that oligodendrocytes could potentially generate a relatively higher proportion of longer Aβ species, which are more prone to aggregation^27^ and, consequently, could be more pathogenic than shorter Aβ species. We therefore used a single-molecule pull-down (SiMPull) assay with DNA-PAINT super-resolution imaging^28^ to analyse soluble Aβ aggregates, of a size consistent with oligomers and protofibrils, in the media from these cells (**Fig. 3e; Supplementary Fig. 7a-b**). This technique relies on dual binding of a single epitope antibody and clustering analysis requiring close proximity of at least two of these super-resolved assemblies to ensure specific detection of soluble aggregates, and has previously been validated to detect stable soluble aggregates with a high sensitivity^28–30^ (see Methods for details). Intriguingly, our analysis revealed that oligodendrocytes produced a greater quantity of these soluble Aβ aggregates than neurons, exceeding the levels that would be anticipated based on the increased amount of Aβ produced (**Fig. 3e-f**). Furthermore, we found that the majority of aggregates produced by both oligodendrocytes and neurons were between 20nm and 200nm in length (**Fig. 3g**; **Supplementary Fig. 7c**), consistent with the reported size of synaptotoxic Aβ aggregates from human AD brains^31^. These results suggest that Aβ produced by oligodendrocytes has a higher propensity for aggregation than neuron-derived Aβ.

### Oligodendrocyte-derived Aβ contributes to plaque formation *in vivo*

To directly investigate whether these features of oligodendrocyte-produced Aβ are consequential for Aβ plaque formation *in vivo*, we generated *App*^NL-G-F^ knock-in mice^32^ with BACE1 knocked out specifically in either oligodendrocytes (BACE1^fl/fl^;PLP1-Cre/ERT^+/-^;*App*^NL-G-F^) or neurons (BACE1^fl/fl^;Thy1-Cre/ERT2,-EYFP^+/-^;*App*^NL-G-F^)., We elected to use the *App*^NL-G-F^ knock-in mouse model for this study as it expresses *App* under its endogenous promoter in a physiological manner with regards to both cell type and amount, and avoids the issue of APP overexpression and abnormal expression patterns. To avoid developmental and early postnatal effects of BACE1 knockout on myelin^33^, we used tamoxifen inducible Cre lines, with tamoxifen administered between the ages of 4 and 8 weeks (after the majority of developmental myelination has taken place^34^), according to standard protocols^35^. Indeed, several control experiments confirmed that this approach had no significant effect on the amount of MBP (**Supplementary Fig. 8a-b**), myelin sheath lengths (**Supplementary Fig. 8c-d**), the number of oligodendrocytes (**Supplementary Fig. 9a-c**) or oligodendrocyte autophagy (**Supplementary Fig. 10a-f**).

We assessed Aβ plaque load within the visual, retrosplenial and motor cortex of these mice at 4 months of age, when there is already widespread plaque distribution within the cortex of *App*^NL-G-F^ mice^32^. We found that oligodendrocyte-specific knockout of BACE1 led to a ∼25% reduction in the number of plaques across the combined area of these cortical regions compared to unmodified *App*^NL-G-F^ control mice (**Fig. 4a-b**), while knockout of BACE1 specifically in neurons led to a near elimination of plaques within the cortex (**Fig. 4a-c**). Notably, oligodendrocyte-specific knockout of BACE1 led to a greater reduction in plaque area in cortical Layers 5/6 as compared to Layers 2/3 specifically within the retrosplenial cortex, which is known to be a selectively vulnerable site of early plaque deposition and functional impairment in AD^36^ (**Supplementary Fig. 11a, e**), albeit not within the motor cortex (**Supplementary Fig. 11b, f**). A similar trend in 20-30% plaque reduction was also observed in the CA1 area of the hippocampus and the corpus callosum, a major white matter tract (**Supplementary Fig. 11c-d, g-h**). These results are consistent with our analysis of snRNA-seq data, where oligodendrocytes had high expression of genes required to produce Aβ in all of the datasets from different brain regions (**Fig. 1a-d**).

**Fig. 4:**
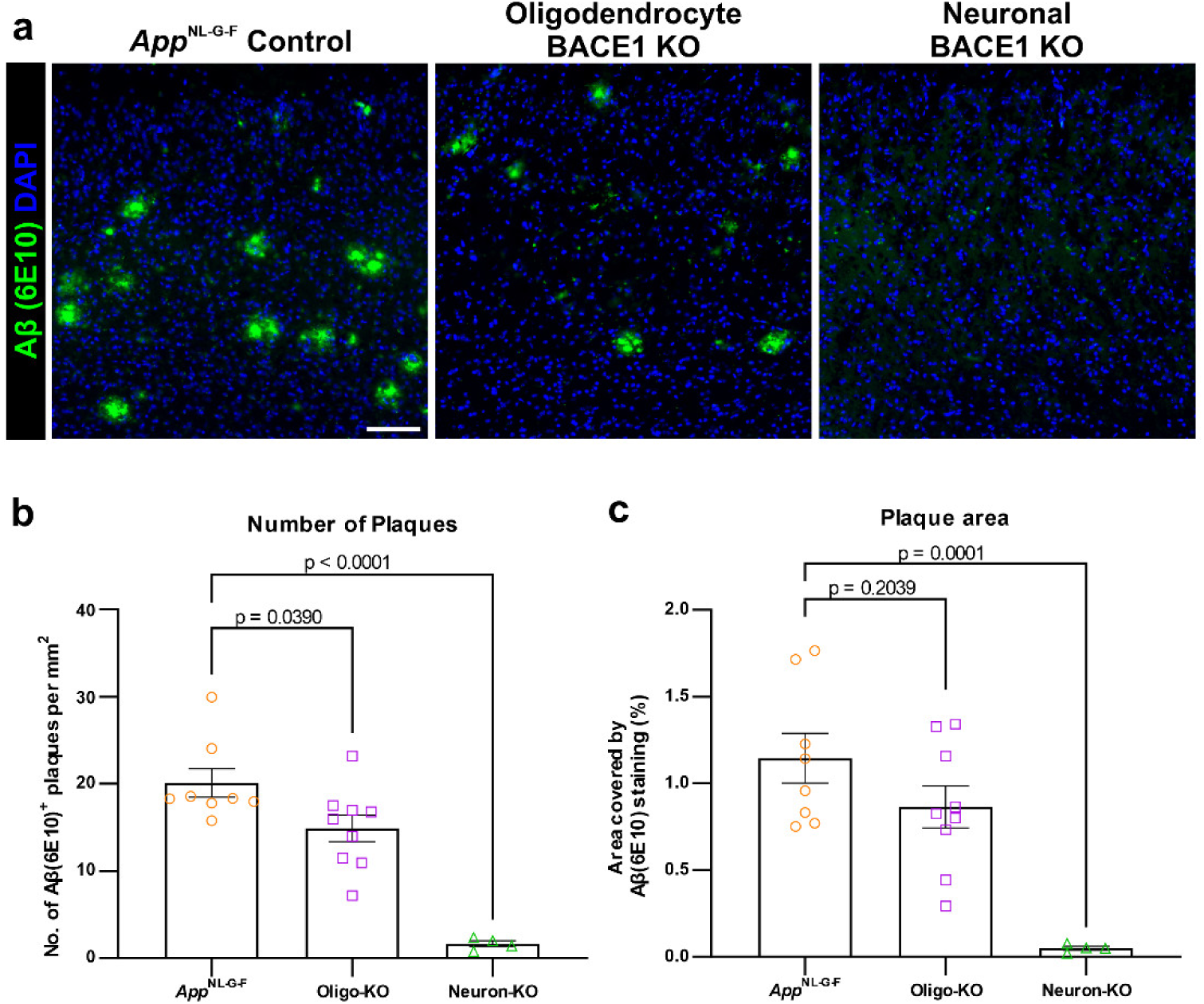
Genetic suppression of oligodendrocyte Aβ production reduces Aβ plaques in the *App*^NL-G-F^ mouse model of AD. **a**, Immunofluorescent images showing Aβ (6E10 antibody; green) and DAPI in the retrosplenial cortex of *App*^NL-G-F^ control mice (left), *APP*^NL-G-F^ mice with BACE1 knocked out (KO) specifically in oligodendrocytes (middle; Oligo-KO), and *APP*^NL-G-F^ mice with BACE1 KO specifically in neurons (right; Neuron-KO). Images show all layers of the cortex, with Layer 6 at the bottom. Scale bar = 100μm. **b,** Quantification of the number of Aβ^+^ plaques across the visual, retrosplenial and motor cortical areas, showing a 25% reduction in Oligo-KO mice compared to *App*^NL-G-F^ and an elimination of plaques in Neuron-KO mice. **c,** Quantification of the total area of Aβ^+^ plaques across the visual, retrosplenial and somatomotor cortical areas, showing a ∼25% reduction in Oligo-KO mice compared to *App*^NL-G-F^ and an elimination of plaques in Neuron-KO mice. In **b,c,** data points represent individual mice (n=8 *App*^NL-G-F^, 9 Oligo-KO, 4 Neuron-KO) with bars showing mean ± SEM. One-way ANOVA with Dunnet’s post-hoc tests: *F*(2,18) = 25.38(**b**), 13.24(**c**); *p*<0.0001(**b**), *p*=0.0003(**c**).

### Suppression of oligodendrocyte-derived Aβ rescues neuronal dysfunction in *App*^NL-G-F^ mice *in vivo*

To determine the consequences of BACE1 knockout in oligodendrocytes on neuronal dysfunction *in vivo*, we used high-density Neuropixels probes to record ongoing neuronal action potential firing in the retrosplenial cortex of 3-month-old awake mice. We found that that oligodendrocyte-specific knockout of BACE1 abolished the early abnormal neuronal hyperactivity phenotype that is present in the *App*^NL-G-F^ mice, which has been shown to be dependent on soluble Aβ^37–39^ (**Fig. 5a-b**). Notably, in *App*^NL-G-F^ mice with an oligodendrocyte-specific knockout of BACE1, we observed that not only were the levels of neuronal firing restored to those seen in WT controls, but so too also the temporal structure, as indicated by the analysis of the variability in mean firing rates and inter-spike intervals (**Supplementary Fig. 12**). We also analysed the power spectrum of cortical local field potentials (LFPs) in the BACE1 knockout mice but found no significant changes compared to controls (**Supplementary Fig. 13**). In addition, analyses of multi-unit neuronal responses in retrosplenial cortex to sharp-wave ripple (SWR) events in CA1 during awake rest^40^ did not reveal any notable impairments in the timing or magnitude of cortical neuronal responses in the BACE1 knockout animals compared to controls (**Supplementary Fig. 14**). Together, these findings indicate that BACE1 knockout in oligodendrocytes effectively ameliorates abnormal neuronal hyperactivity in *App*^NL-G-F^ mice without disrupting the electrophysiological integrity of the cortical-hippocampal network.

**Fig. 5:**
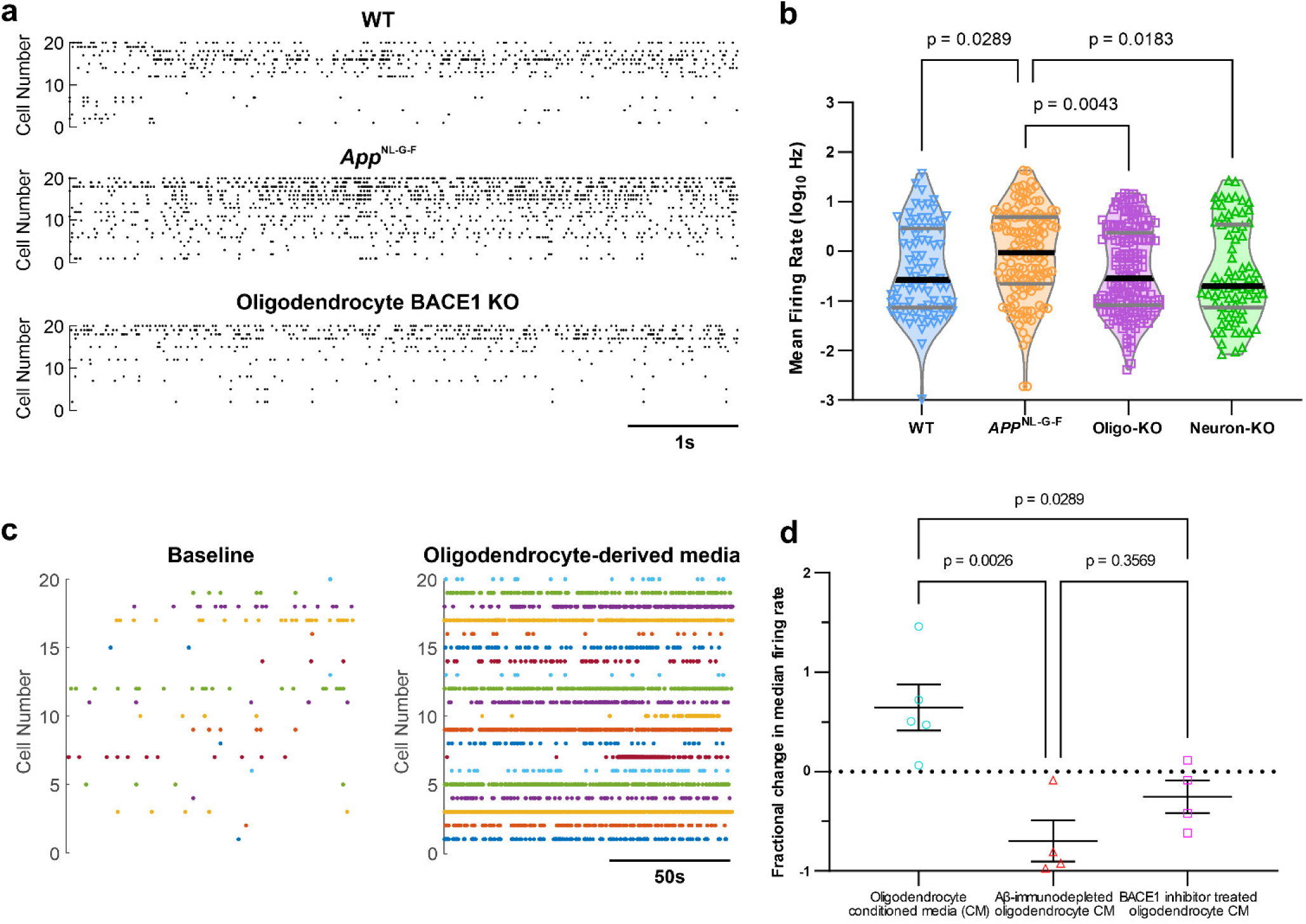
Genetic suppression of oligodendrocyte Aβ production rescues neuronal dysfunction in the *App*^NL-G-F^ mouse model of AD, while oligodendrocyte-derived media promotes neuronal dysfunction *in vivo*. **a**, Raster plots from Neuropixels recordings showing spontaneous neuronal firing in 20 randomly selected cortical neurons/units from 3-month-old awake WT (top), *App*^NL-G-F^ (middle) and oligodendrocyte BACE1 KO (bottom) mice, illustrating rescue of hyperactivity phenotype in oligodendrocyte BACE1 KO mice to WT levels. Units are sorted from high mean firing rates to low (top to bottom) and plots show a 6s resting state period. **b,** Quantification of the mean firing rate showing significantly reduced activity in cortical neurons of both Oligo-KO and Neuron-KO mice compared to *App*^NL-G-F^ mice, with firing rates returning to levels observed in WT controls (WT vs. Oligo-KO: *p* > 0.9999; WT vs. Neuron-KO: *p* > 0.9999). In **b,** data points represent individual neurons/units (n=82 WT, 134 *App*^NL-G-F^, 165 Oligo-KO, 75 Neuron-KO) across 4 (WT) or 3 mice per group with the shaded area representing smoothed distribution. Median is shown by a thick black line, and quartiles indicated with grey lines. Kruskal-Wallis test (*H*(3, n=456)=15.33, *p*=0.0016) with Dunn’s post-hoc tests. **c,** Raster plots showing neuronal firing in the same 20 cortical neurons/units at baseline (left) and during injection of oligodendrocyte conditioned media containing soluble Aβ aggregates (right) in 4-month-old WT mice, illustrating the strong increase in neuronal activity upon exposure to oligodendrocyte-derived Aβ. **d,** Quantification showing an increase in neuronal firing rates upon local injection of oligodendrocyte conditioned media (Aβ_40_ concentration by ELISA: 133 pM) into retrosplenial cortex, compared to injection of either the same media which has been immunodepleted of Aβ (Aβ_40_ concentration by ELISA: 24 pM) or media from BACE1 inhibitor-treated oligodendrocytes (Aβ_40_ concentration by ELISA: 18 pM). In **d,** data points represent individual mice (n=5 oligodendrocyte conditioned media, 4 other groups) with mean ± SEM shown. One-way ANOVA (*F*(2,10)=11.23, *p*=0.0028) with Tukey’s post-hoc tests.

Our results that targeted suppression of Aβ production in oligodendrocytes can rescue neuronal hyperactivity, even without the complete elimination of plaques, indicated that soluble Aβ species produced by oligodendrocytes may in fact contribute to early neuronal dysfunction in the AD brain. To directly test this hypothesis, we injected human fAD iPSC-derived oligodendrocyte conditioned media containing soluble Aβ aggregates into the retrosplenial cortex of WT mice while recording ongoing neuronal action potential firing using Neuropixels *in vivo*. As controls, we administered the same media immunodepleted of Aβ using 6E10 antibody, or media from oligodendrocytes which had been treated with BACE1 inhibitor in order to suppress Aβ production. Indeed, we found that neuronal firing rates were markedly increased during injection of the oligodendrocyte conditioned media compared to baseline, which was not observed with control media (**Fig. 5c-d**). These results suggest that oligodendrocyte-derived Aβ is sufficient to promote neuronal hyperactivity, in the absence of Aβ from any other cellular source.

## Discussion

In conclusion, we have shown that human oligodendrocytes not only produce Aβ, but they can also generate Aβ in greater amounts and with a higher proportion as soluble aggregates compared to neurons. This was seen in oligodendrocytes both with and without fAD mutations. The increased Aβ 42:40 ratio in oligodendrocytes could contribute to the high proportion of soluble aggregates^27,41,42^; it is also worth noting that the relationship between Aβ monomer concentration and aggregate concentration is non-linear, so small changes in monomer can result in larger increases in the number of aggregates^43^, although this does not rule out additional contributing factors. We demonstrate that specific suppression of oligodendrocyte Aβ production is sufficient to rescue neuronal hyperactivity in the *App*^NL-G-F^ knock-in model of AD. In turn, administration of oligodendrocyte conditioned media containing soluble Aβ promotes neuronal hyperactivity in WT mice *in vivo*. This result is consistent with and extends previous studies on the effects of soluble Aβ on neuronal activity, although these earlier studies did not consider the cellular source of Aβ^37–39^.

The functional rescue is remarkable given the relatively modest reduction in plaque load that results from blocking oligodendrocyte Aβ production, while blocking neuronal Aβ production leads to a near elimination of plaques. This aligns with work from multiple laboratories showing that plaque formation depends on neuronal Aβ ^44–47^, and is consistent with the lower abundance of plaques observed in the white matter compared to the grey matter, although elevated levels of soluble Aβ have previously been reported in the white matter of individuals with AD^48^. The lower abundance of plaques in the white matter could further be explained by the evidence that most neuronal Aβ is secreted at presynaptic terminals, thus favouring plaque deposition in the grey matter^49,50^. This small contribution of oligodendrocytes to plaque load could suggest that a main effect of oligodendrocyte-derived Aβ is to promote neuronal dysfunction. Rather than being plaque-dependent, the effects of Aβ on neuronal activity and synaptic function often stem from soluble aggregates similar to those we see produced in much greater numbers by oligodendrocytes^31,38^.

Together with our data showing an increased number of Aβ-producing oligodendrocytes in deeper cortical layers of the brains of individuals with AD, these results indicate that oligodendrocyte-derived Aβ plays a pivotal role in the early impairment of neuronal circuits in AD^51^, which has important implications for how we consider and treat the disease. The increased number of oligodendrocytes in human AD brains also raises the intriguing possibility that these cells could potentially offset reduced Aβ production due to neuronal loss as the disease progresses. While recent anti-Aβ antibody therapies targeting plaques have been shown to be effective in slowing the clinical course of AD, they can lead to blood vessel damage, detected as amyloid-related imaging abnormalities (ARIA) on MRI^52^. Previous trials to reduce Aβ through BACE inhibition have failed to demonstrate similar beneficial effects, which is thought to be due to the effects on other neuronal BACE substrates such as SEZ6^53^, which, importantly, is not expressed in oligodendrocytes (**Supplementary Fig. 15**). Thus, to circumvent such unwanted effects, we propose that blocking Aβ production specifically in oligodendrocytes, for example by employing oligodendrocyte-targeting AAVs^54^, may constitute a promising novel target for treating AD.

## Acknowledgements

We thank David Attwell, Siddharthan Chandran, Bart De Strooper and James Rowland for comments on the manuscript. We thank UK DRI at UCL technical staff Elena Ghirardello and Phillip Muckett for the maintenance of animal colonies and assistance with animal experiments. We thank the Queen Square Brain Bank for Neurological Disorders for provision of human brain tissue samples. The Queen Square Brain Bank is supported by the Reta Lila Weston Institute of Neurological Studies, UCL Queen Square Institute of Neurology. We thank Tanja Kuhlmann and the University of Münster for providing the SON lentivirus. We thank Takashi Saito and Takaomi C. Saido for provision of *App*^NL-G-F^ mice. We thank Ulf Neumann and Derya Shimshek from Novartis for providing us with the BACE1 inhibitor NB-360. This work was supported by the UK Dementia Research Institute through UK DRI Ltd, principally funded by the UK Medical Research Council. This work was further supported by the National Institute for Health Care Research University College London Hospitals Biomedical Research Centre. S.S.H. is further supported by an Alzheimer’s Association International Research Fellowship (AARF-23-1149637). F.K.W.L. is funded by an NC3R studentship (NC/W001675/1). M.S. is supported by an MRC Career Development Award (MR/X019977/1). T.A.G. is supported by an Alzheimer’s Association Research Fellowship to Promote Diversity (23AARFD-1029918). M.A.B. is supported by an UKRI Future Leaders Fellowship (MR/X011038/1).

## Author contributions

R.M.R. conceptualised the study, and designed, carried out and analysed most of the experiments. R.E. carried out Neuropixels experiments and processed the data. M.H. and O.S.T. carried out *in vitro* experiments and ELISAs, and O.S.T. carried out ISH experiments. S.S.H constructed the Neuropixels rig and analysed the Neuropixels data. D.G. and F.K.W.L assisted with *in vitro* experiments and ELISAs, and D.G. carried out IHC experiments, western blots and qPCR. C.A., T.A.G. and S.W. assisted with *in vitro* experiments and provided iPSCs. E.F., J.S.H.D. and D.K. carried out imaging and analysis of Aβ aggregates. M.S. and M.L. carried out IHC, imaging and analysis of myelin sheath lengths. M.B. carried out IHC and analysis of autophagy. R.V. provided BACE1^flox^ mice. C.S.F. initiated the original experiments, carried out snRNA-seq analysis, contributed to study conceptualisation and design, and provided supervision. M.A.B. contributed to study conceptualisation and design as well as data interpretation. R.M.R. and M.A.B. wrote the manuscript with input from all authors. M.A.B. supervised all aspects of the project.

## Competing interests

The authors declare the following competing interests: C.S.F. is currently employed by GSK.

## Data availability

The full dataset is available from the corresponding authors upon request and will be stored on UCL Research Data Repository: (https://www.ucl.ac.uk/library/openscience-research-support/research-data-management/ucl-research-data-repository). All raw data necessary to reproduce all figures will be available within the Source data provided with this paper.

## Methods

### Animals

All experimental procedures were conducted in accordance with the Animal (Scientific procedures) Act 1986, approved by the Animal Welfare and Ethical Review Body at University College London (UCL), and performed under an approved UK Home Office project licence at UCL. Mice were maintained in a 12-hour light/dark cycle with food and water supplied *ad libitum*. *App*^NL-G-F^ mice [C57BL/6-App<tm3(NL-G-F)Tcs> (No.RBRC06344)] were provided by the RIKEN BRC through the National BioResource Project of the MEXT, Japan^1^. BACE1^fl/fl^ mice (C57BL/6– Bace1tm1.1mrl) were previously generated and provided by R.V.^2^. PLP1-Cre/ERT mice [B6.Cg-Tg(Plp1-cre/ERT)3Pop/J; JAX:005975] and Thy1-Cre/ERT2,-EYFP mice [B6.Tg(Thy1-cre/ERT2,-EYFP)HGfng/PyngJ; JAX:012708] were purchased from The Jackson Laboratory. Three mouse cohorts were generated: (1) BACE1^fl/fl^; PLP-Cre/ERT^+/-^;*App*^NL-G-F^, (2) BACE1^fl/fl^; Thy1-Cre/ERT,-EYFP^+/-^;*App*^NL-G-F^, and (3) BACE1^fl/fl^; *App*^NL-G-F^ (i.e., the Cre/ERT^-/-^ littermates from the other two cohorts). All three cohorts were treated with tamoxifen between 4 and 8 weeks of age: tamoxifen (MP Biomedicals; 156738) was dissolved in corn oil at 25 mg/ml. Animals were injected with tamoxifen intraperitoneally at a dose of 100 mg/kg daily for 5 days followed by a 2 day break, repeated 4 times. For Neuropixels recordings, C57BL/6J mice (originally purchased from Charles River Laboratories) were used as wild type (WT) controls. For acute recordings, animals were aged between 4 and 5 months. All other animals were aged between 3 and 4 months at the time of Neuropixels recordings or perfusion for fixed tissue analyses. Data presented shows the means of both male and female animals.

### Immunohistochemistry (IHC)

Brains were extracted from animals after intracardiac perfusion of 1x phosphate buffered saline (no calcium no magnesium; PBS; Gibco), post-fixed for 24 hours in 4% paraformaldehyde (PFA) in PBS (Alfa Aesar), cryopreserved through increasing sucrose concentrations, embedded in optimal cutting temperature compound (OCT; CellPath) and frozen on dry ice. Amyloid plaque and oligodendrocyte analyses: 10 μm thick sagittal sections were cut using a Leica cryostat. Sections were blocked in blocking solution (10% donkey or goat serum, 0.3% triton in PBS) for 3 hours then incubated with Alexa Fluor 594 anti-β-amyloid (Clone 6E10; 1:1,000; BioLegend; 803019), Olig2 (1:500; Millipore; MABN50), ASPA (1:1,000; Sigma-Aldrich; ABN1698), APP (Y188; 1:500; Abcam; ab32136), and/or BACE1 (1:250; Abcam; ab108394) antibodies in block overnight at 4°C. Where appropriate, sections were washed and incubated with secondary antibodies [Goat anti-Mouse Alexa Fluor 488 (1:1,000; Invitrogen; A11001), Goat anti-Rabbit Alexa Fluor 594 (1:1,000; Invitrogen; A11012), Goat anti-Mouse Alexa Fluor 594 (1:1,000; Invitrogen; A11005), Goat anti-Rabbit Alexa Fluor 488 (1:1,000; Invitrogen; A11008)] in block for 2 hours at room temperature. Sections were then labelled with DAPI and mounted with ProLong Gold. Myelin sheath analysis: 30μm thick coronal brain sections were cut between bregma +1.1mm and +0.8mm. Free-floating coronal sections were blocked in 10% fetal calf serum, 0.1% triton in PBS for 1 hour then incubated with rat anti-MBP antibody (1:500; Abcam; ab7349) in blocking solution overnight at 4ᵒC. Sections were washed in PBS before adding Donkey anti-Rat Alexa Fluor 488 secondary antibody (1:1000; Invitrogen; A21208) for 1 hour at room temperature. Sections were washed and stained for nuclei before being mounted onto slides. Oligodendrocyte autophagy analysis: 40μm thick free-floating coronal sections were washed in PBS, blocked in M.O.M. blocking solution (Vector laboratories) for 90 minutes, then incubated in primary antibodies [Olig2 (1:500; Millipore; MABN50), LC3 (1:1,000; Sigma-Aldrich; L7543), Lamp2 (1:2,000; Hybridoma Bank; ABL-93), p62 (1:500; Abcam; ab91526), Ctsd (ThermoFisher; PA5-47046)] overnight at 4°C. Sections were then washed in PBS followed by incubation with secondary antibodies (Donkey anti-Mouse 488, Donkey anti-Mouse 568, Donkey anti-Rabbit 657, Donkey anti-Rat 488, Donkey anti-Goat 568; all at 1:1,000; Invitrogen) for 2 hours at room temperature. Sections were then washed, labelled with DAPI and mounted onto slides.

### Western blotting

Brains were extracted from animals after intracardiac perfusion of 1x PBS, the forebrain was dissected and protein extracted in RIPA buffer (Pierce). Protein in each sample was quantified using a BCA Assay (Pierce) and 20μg of protein was loaded per well of a NuPAGE 4-12% Bis-Tris gel. Samples were run for 1.5 hours at 150V, before transfer to a nitrocellulose membrane using an iBlot 2 transfer system (ThermoFisher Scientific). Membranes were blocked for 1 hour in 1% casein (Bio-Rad) in PBST (0.1% Tween-20 in PBS), then incubated with MBP (1:1,000; Bio-Rad; MCA409S) antibody overnight at 4°C. Membranes were washed in PBST, incubated with Goat anti-rat ECL antibody (1:10,000; Amersham; NA935) for 1 hour, then imaged using Pierce ECL substrate and an Amersham Imager 680. Membranes were subsequently stripped (Pierce) and reprobed for GAPDH [(1:1,000; Sigma-Aldrich; MAB374); secondary: Sheep anti-mouse ECL (1:10,000; Amersham; NA931)]. Blots were analysed using the Gel Analyzer tools in the FIJI build of ImageJ (open source). Intensity of MBP bands was normalised to the GAPDH band for each lane, and all lanes were normalised to the average of the *App*^NL-G-F^ values.

### Neuropixels recordings and analysis

#### Surgical Procedures – Awake recordings

General anaesthesia was induced using ∼4% isoflurane in O_2_ and maintained at ∼2% throughout the surgery, with anaesthetic depth monitored via the pedal reflex and breathing rate. Carprofen for pain relief was administered by subcutaneous injection prior to surgical procedures. Animals were head fixed in a Mouse Ultra Precise Stereotaxic Instrument (World Precision Instruments, WPI) using ear bars. Normal body temperature was maintained using a heating pad and Viscotears gel applied to the eyes. The scalp was shaved, disinfected with dilute chlorhexidine and cleaned with alcohol, and treated for 5 minutes with lidocaine cream before skin excision. Following exposure, the skull was first covered with a thin layer of Vetbond (3M), followed by ultraviolet curing Optical Adhesive 81 (Norland), which was cured using a handheld LED UV spot lamp (Intertronics). A craniotomy spanning approximately half the area of the left and right parietal bones between bregma and lambda was drilled using an OmniDrill35 (WPI). A titanium headplate was subsequently attached to the skull behind lambda with SuperBond dental cement (Prestige Dental) and a circular well encircling the exposed skull created using dental cement (Jet, Lang). A layer of sterile PBS was applied to the craniotomy before sealing the well with KwikCast silicone elastomer (WPI). Animals were taken off isoflurane, received a subcutaneous buprenorphine injection for immediate pain relief and were provided with drinking water containing carprofen for 3 days postoperatively. Body weight and score were monitored to ensure appropriate recovery and health of the animal.

#### Surgical Procedures – Acute recordings

Surgical procedures for animals undergoing subsequent acute Neuropixel recordings under isoflurane anaesthesia were conducted as above, but forewent ultraviolet curing and headplate attachment and underwent additional insertion of a silver chloride reference electrode into a small hole drilled over the cerebellum and secured with cyanoacrylate glue and dental cement. These animals were transferred directly to the recording stage at the end of surgical procedures and without interruption of isoflurane anaesthesia.

#### Recording Procedures – Awake

Two weeks post-surgery, mice were put through three daily habituation sessions to familiarise them to handling procedures, head fixation apparatus and Neuropixels recording setup. Animals were secured via the implanted headplate in a holder (Thorlabs) suspended above a fixed running wheel fitted with a rotary encoder to monitor locomotion. Behaviour was also monitored via an IR camera filming the head and forelimbs of the animal during Neuropixels recordings. Mice were then lightly sedated with <1% isoflurane, the silicone cap removed, and a silver chloride reference electrode placed in contact with the skull and secured to the edge of the well using dental cement (Jet, Lang). A Neuropixels probe (IMEC) was secured to a QUAD micromanipulator (Sutter Instruments) orthogonal to the anterior posterior axis of the mouse at a 60° elevation angle and spatially referenced to bregma. The probe was then maneuvered to 2.2 mm posterior and 0.5 mm medial of bregma so as to be inserted into the right hemisphere along a trajectory spanning the retrosplenial cortex and CA1 region of the hippocampus. The probe was then inserted at ∼10 µm/s along the elevation angle to a depth of 3.3 mm. Following implantation, anaesthesia was removed. 10 min recordings of spontaneous brain activity sampled at ∼30 kHz were taken 40 minutes after probe implantation. After recordings, the probe was slowly retracted, the reference electrode detached, and well refilled with sterile PBS and KwikCast before animals were returned to their home cage.

#### Recording Procedures – Acute media injections

Following surgical procedures, mice were transferred to the recording stage without interruption of anaesthesia and while remaining in the stereotaxic frame. A 10µl Hamilton needle syringe, secured to a microinjection pump (UMP3, WPI), was then carefully inserted into retrosplenial cortex (∼2mm AP, 0.5mm ML) to a depth of ∼600µm under micromanipulator control (MM33, Multichannel Systems) and visualised under a microscope (GT Vision). The syringe was preloaded with control media (media conditioned by BACE1 inhibitor-treated oligodendrocytes, or media which had been immunodepleted of Aβ; see details below) or oligodendrocyte conditioned media (media conditioned by vehicle-treated oligodendrocytes grown in parallel, containing 400-500nM of Aβ aggregates, as estimated by aggregate quantification, and 130pM of Aβ_40_ monomers, as measured by ELISA; see details of media collection and quantification below). Once in position, a Neuropixels probe (IMEC) was secured to a QUAD micromanipulator (Sutter Instruments) and implanted as for awake recordings and as close as possible to the injection needle. Following implantation, the brain was allowed to rest for at least 40 minutes before subsequent recordings, during which isoflurane anaesthesia was gradually lowered to maintain adequate anaesthetic depth and promote physiological cortical activity (∼1%). A single recording of spontaneous brain activity was then performed during which, following a baseline acquisition period of 5mins, media was injected at a rate of 10nl/min over 10 mins for a total injection volume of 100nl into retrosplenial cortex.

#### Neuropixels Analysis

Action potential (i.e. spiking) data (sampling frequency 30kHz) were separated into individual units using the automated spike sorting algorithm Kilosort3^3^. For awake recordings, averaged spike waveforms associated with each identified unit were extracted and normalised, and the first and second principal components computed for each waveform (Matlab function ‘pca’). These metrics were used as inputs to generate a Gaussian mixture distribution model (GMM, Matlab function ‘fitgmdist’, 200 replicates) for subsequent clustering with two components (Matlab function ‘cluster’), with the smallest cluster (typically accounting for <10% of all unit waveforms and consisting largely of artefactual signals) excluded from further analysis. Remaining waveforms were subsequently visually checked to confirm physiological appearance.

To map identified units to anatomical brain regions we used custom-written Matlab scripts comprising a modification to the open-source Neuropixels trajectory explorer (https://github.com/petersaj/neuropixels_trajectory_explorer) which incorporates the Allen Common Coordinate Framework (CCF) v3 mouse atlas. This produced a putative anatomical trajectory of the probe with respect to the stereotactic implant coordinates and as a function of the recording channel along the probe shank. To offset this trajectory to account for implantation depth, a depth template of probe channels was created, where cortical and CA1 regions in the predicted trajectory were assigned a non-zero value, with all other areas set to zero. This created a template consisting of two square waves that reflects the span of the cortex and CA1 separated by the intervening corpus callosum. The estimated offset was calculated as the largest cross-correlation of this template with a profile of the computed sum of multi-unit spiking activity across each recording channel (Matlab function ‘finddelay’). This calculated offset was then used to align the anatomical trajectory to functional data, so as to assign isolated units to putative brain regions, and visually confirmed.

Behavioural data collected from the IR camera video and rotary encoder were used to separate recordings into different behavioural states. Stationary/quiescent periods were defined as timepoints where angular acceleration on the rotary encoder was < 0.025 absolute rotations per minute (RPM)/sec and velocity was < 0.01 RPM. Non-locomotor movements, such as grooming, whisking etc., were identified by first calculating the rectified first approximate derivative of the mean intensity value across all pixels in each video frame and defined as signals crossing above 6-fold median absolute deviations. Behavioural data were then split into multiple 1 second windows uninterrupted by locomotion or non-locomotor movements. Windows in which the theta (4-8Hz) – delta (< 4Hz) power ratio remained less than 2 (calculated using the mean local field potential across cortical channels, Matlab function ‘pspectrum’, 1s time resolution with 99% overlap)^4^ were subsequently defined as resting state epochs. Spontaneous mean firing rate (MFR) was calculated for each unit over all identified resting state epochs in each animal across 2-3 concatenated Neuropixels recording sessions (mean total duration ∼28min). We additionally calculated, for each neuron/unit, the coefficient of variation (CV, standard deviation divided by the mean) of the MFR across resting state epochs (“CV MFR”), as well as the coefficient of variation 2^5^ of the inter-spike interval (ISI) during quiescent (i.e. non-locomotion) states (“CV2 ISI”).

For acute experiments involving local media injections, we calculated the firing rate of all identified units over 1s bins during the recording period and averaged over the 120s baseline period prior to media injection, and that during the last 120s of media injection, to obtain baseline and end-of-injection mean FRs. For each experiment (i.e. animal), the median firing rate across all units was calculated for both baseline and end-of-injection conditions, and the fractional change in median firing rate calculated as (medianFR*_end-of-injection_* – medianFR*_baseline_*) / medianFR*_baseline_*.

#### Local field potential analysis

Broadband local field potential data (LFP, sampling frequency 2.5kHz) were selected from dorsal retrosplenial cortex-associated channels and common average referenced. For each such channel, stationary/quiescent periods were sub-divided into complete 60s epochs and power spectral density estimates for each epoch successively calculated using the multitaper method (Matlab function ‘pmtm’, 7 Discrete Prolate Spheroidal (Slepian) sequences as tapers, <100Hz), and averaged within and then across recording sessions for each animal. Overall power was subsequently calculated by integrating the associated area using the Simpson’s rule (parabolic approximation).

#### Sharp wave ripple (SWR) analysis

Local field potential data from CA1 channels were common average referenced and filtered for the ripple band (110 to 250 Hz; 4th order Butterworth filter with Second-Order-Section implementation for stability), and envelopes/instantaneous amplitudes calculated using the Hilbert transform (Matlab functions ‘hilbert’ and ‘abs’). SWR events were detected similarly to previously published protocols^6^. Local maxima in the ripple band envelope which exceeded five times the median of the envelope values within that channel during quiescence were collated as candidate SWR events. Onset and offset for each event were defined as the nearest timepoints prior to and following local maxima in which ripple band envelope fell below 50% of the detection threshold, with SWR event time defined as the time of the local maxima. Candidate events were included for final analysis if ripple band power during the event was at least twice as large as that in the common average reference and that of the power in the supra-ripple band (200-500Hz, to account for high frequency noise), occurred when the animal was quiescent and if the theta/delta power ratio was <2 (i.e. during the resting state), and if the event was at least 4 ripple cycles in length and separated from another by at least 500 ms. Similarly to previous protocols^7,8^, we next examined the multi-unit cortical response to each ripple by computing the ripple triggered peri-event time histograms (PETHs) and counting total spiking activity across all identified dorsal retrosplenial neurons/units in 10ms bins (range –1 to +1s). PETHs were smoothed with a 50 ms Gaussian filter, averaged over SWR events across experimental sessions for each animal (minimum of 100 events for inclusion) and z-scored.

### Human tissue

Post-mortem human brain tissue from prefrontal cortex (Brodmann’s Area 10) was provided as pre-cut frozen tissue sections by the Queen Square Brain Bank with ethical committee approval (reference UCLMTA4-20), and the tissue is stored under licence from the Human Tissue Authority (licence number 12198). Alzheimer’s Disease (AD) tissue is from donors with a clinical diagnosis of sporadic AD and a high level of AD pathological change (A3, B3, C2 or C3). Control tissue is from donors with no clinical diagnosis of neurological disease and no or minimal AD-related pathology. Age: control, 90 ± 7 years (mean ± SD) and AD, 75 ±10; Post-mortem delay: control, 69 ± 25 hours and AD, 59 ± 28; Percentage male: control, 50% and AD, 80%.

### RNAscope *in situ* hybridization (ISH)

ISH was carried out using RNAscope Multiplex Fluorescent Kit v2 [Advanced Cell Diagnostics (ACD)] according to the manufacturer’s instructions. Briefly, frozen 10 µm thick human tissue sections were dried [4 minutes at 40°C then 10 minutes at room temperature (RT)] before fixation in 4% PFA in PBS (30 minutes at RT). Sections were then dehydrated through increasing ethanol concentrations (5 minutes each in 50%, 70%, 100%, 100% ethanol) before incubation with hydrogen peroxide (10 minutes at RT) followed by incubation with Protease IV (20 minutes at RT). Hybridization of the probes [Hs-APP (418321; ACD), Hs-BACE1-C2 (422541-C2; ACD), and Hs-MBP-C3 (411051-C3; ACD) or Hs-RBFOX3-C3 (415591-C3; ACD) at a ratio of 50:1:1] was carried out at 40°C for 2 hours. Sections then underwent amplification (30 minutes at 40°C for each probe sequentially) followed by sequential detection of each probe [15 minutes at 40°C with probe-specific HRP then 30 minutes at 40°C with Opal dye (1:500; C1:Opal 570, C2: Opal 620, C3: Opal 520; Akoya Biosciences)]. Following ISH, sections underwent IHC for Aβ: sections were blocked for 1 hour in blocking solution (10% goat serum, 0.1% triton in PBS), incubated with primary antibody in block (anti-β-amyloid clone 6E10; 1:2,000; BioLegend; 803001) overnight at 4°C, then incubated with secondary antibody in block [Goat anti-Mouse Alexa Fluor 647 (1:1,000; Invitrogen; A21235)] for 2 hours at RT. Sections were incubated with TrueBlack (1X; Biotium; 23007) for 30 seconds and counterstained with DAPI (1:5,000) prior to mounting with ProLong Gold (Invitrogen).

### Induced pluripotent stem cells (iPSCs)

iPSCs were previously obtained or generated, and validated, by C.A. and S.W.^9,10^ iPSCs were cultured in Essential 8 media on Geltrex and passaged using 0.5mM EDTA (all reagents from ThermoFisher). The following cell lines were used in this study: APP V717I 1 (APP V717I mutation), PSEN1 R278I (PSEN1 R278I mutation), PSEN1 int4del (intron 4 deletion in PSEN1), PSEN1 WT (PSEN1 int4del line CRISPR edited to correct the mutation^9^).

### iPSC-derived cortical neurons

Cortical neurons were derived from iPSCs as previously described^11^. Briefly, iPSCs underwent neural induction in N2B27 media [1:1 DMEM-F12 Glutamax (Gibco): Neurobasal (Gibco) with 0.5X B27 supplement (Gibco), 0.5X N2 supplement (Gibco), 0.5X Non-essential amino acids (Gibco), 50 μM 2-mercaptoethanol (Gibco), 2.5 µg/ml insulin (Sigma-Aldrich), 50 U/ml Penicillin (Gibco), 50 µ/ml Streptomycin (Gibco), 1 mM L-glutamine (Gibco)] supplemented with 1 µM Dorsomorphin (Tocris) and 10 µM SB-431542 (Generon) for 10 days. Cells were then expanded over 12-18 days in neural maintenance media, and split every 6 days using dispase (Gibco) to remove non-neuronal cell types. Cells were then singularised using accutase (Sigma-Aldrich) before final plating in poly-L-ornithine (Sigma-Aldrich) and laminin (Sigma-Aldrich) coated wells. Mature cortical neurons begin to appear from 60 days after final plating, and media which had been on the cells for 24 hours was collected for Aβ quantification between 120 and 200 days after final plating. To normalise for cell number, cells were lysed at the end of the experiment, total genomic DNA extracted and quantified, and cell number calculated based on each cell containing 6pg of DNA.

### iPSC-derived oligodendrocytes and OPCs

Oligodendrocytes were derived from iPSCs as previously described^12^. Briefly, neural precursor cells (NPCs) were first derived from iPSCs as previously described^13^: iPSC colonies were grown in N2B27 media supplemented with 1 µM Dorsomorphin (Tocris), 10 µM SB-431542 (Generon), 3 µM CHIR 99021 (MedChemExpress) and 0.5 µM purmorphamine (Peprotech). After 4 days, Dorsomorphin and SB-431542 were withdrawn and 150 µM ascorbic acid was added. Cells were plated on Geltrex (Gibco), and grown for 30 days, with passaging every 5-6 days, to expand and remove contaminating cell types. NPCs were then plated on Geltrex at a density of 100,000 cells/well and expression of *SOX10*, *OLIG2* and *NKX6.2* was driven by transduction of the cells with SON lentivirus (plasmid kindly provided by Prof. Tanja Kuhlmann, University of Münster; virus constructed by VectorBuilder) at an MOI of 5, together with 5 µg/ml protamine sulfate (Sigma-Aldrich) for 24 hours. Cells were then grown in glial induction medium [GIM; DMEM-F12 glutamax with 0.5X B27 minus vitamin A supplement, 0.5X N2 supplement, 100 U/ml Penicillin, 100 µg/ml Streptomycin, 2 mM L-glutamine, 1 µM SAG (Cayman Chemical), 10 ng/ml PDGF (Peprotech), 10 ng/ml NT3 (Peprotech), 10 ng/ml IGF-1 (Peprotech), 200 µM ascorbic acid, 0.1% Trace Elements B (Corning), 10 ng/ml T3 (Sigma-Aldrich)]. After 4 days, cells were changed into differentiation media [DM; DMEM-F12 glutamax with 0.5X B27 minus vitamin A supplement, 0.5X N2 supplement, 100 U/ml Penicillin, 100 µg/ml Streptomycin, 2 mM L-glutamine, 60ng/ml T3, 10ng/ml NT3, 10ng/ml IGF-1, 200 µM ascorbic acid, 0.1% Trace Elements B, 100 µM dbcAMP (Sigma-Aldrich)]. After 7-10 days of differentiation, cells were singularised with accutase and replated on Geltrex at a density of 250,000 cells/well (12-well-plate) or 150,000 cells/well (glass coverslips in 24-well-plate). 35 days after glial induction, almost all cells were oligodendroglia (OLIG2^+^ and/or MBP^+^) and cells with mature MBP^+^ sheets were visible. Media which had been on the cells for 24 hours was collected for Aβ quantification between 40 and 120 days after glial induction. For immature OPCs, cultures were maintained in GIM, and media which had been on the cells for 24 hours was collected for Aβ quantification at day 5 after glial induction. To normalise for cell number, cells were lysed at the end of the experiment, total genomic DNA extracted and quantified, and cell number calculated based on each cell containing 6pg of DNA.

### iPSC-derived astrocytes

iPSC-derived astrocytes were generated using an established protocol^14^. Briefly, glial progenitor cells were enriched from cortical neuronal cultures (see above) in a proliferative phase [N2B27 media plus 10ng/ml FGF2 (Peprotech)] until day 150 of differentiation. During this phase, glial progenitors were routinely passaged using 0.5mM EDTA and cultured on Geltrex substrate (ThermoFisher). A final two-week maturation phase consisted of 10ng/ml LIF (Sigma) and 10ng/ml BMP4 (Peprotech), at the end of which media which had been on the cells for 24 hours was collected for Aβ quantification. To normalise for cell number, cells were lysed at the end of the experiment, total genomic DNA extracted and quantified, and cell number calculated based on each cell containing 6pg of DNA.

### iPSC-derived microglia

iPSC-derived microglial-like cells were generated using established protocols^15,16^. Briefly, embryoid bodies were generated from 10,000 iPSCs and haematopoietic differentiation was initiated via 3 days of 50ng/ml BMP4, 50ng/ml VEGF and 20ng/ml SCF followed by maintenance in myeloid media consisting of x-vivo media with 25ng/ml IL3 and 100ng/ml MCSF. Microglial-like cells were harvested and exposed to a two-week final maturation in 25ng/ml MCSF, 100ng/ml IL34, 5ng/ml TGF-β and further supplemented with CX3CL1 and CD200 for the final 2 days (both 100ng/ml), at the end of which media was collected for Aβ quantification. All growth factors Peprotech unless stated. To normalise for cell number, cells were lysed at the end of the experiment, total genomic DNA extracted and quantified, and cell number calculated based on each cell containing 6pg of DNA.

### BACE1 inhibitor treatment

BACE1 inhibitor NB-360 (a kind gift from Novartis) was prepared at a concentration of 10 µM in differentiation medium (DM), from a stock solution of 10 mM prepared in dimethyl sulfoxide (DMSO; Sigma-Aldrich). Cells between 40 and 60 days after glial induction which had been in their medium for 24 hours had their medium collected (pre-treatment) and medium containing 10 µM NB-360 was added. After 24 hours of treatment, the medium was collected (post-treatment) and replaced with regular DM. As vehicle controls, cells were in parallel treated with DM containing 0.1% DMSO. Aβ concentrations were quantified in both pre-treatment and post-treatment media samples to calculate the change in Aβ production for each well of cells.

### DNA extraction

DNA was extracted using the QIAGEN DNeasy Blood and Tissue Kit (69504) according to the manufacturer’s instructions. Briefly, cells were collected and suspended in PBS with 10% Proteinase K. Lysis buffer was added and the samples were incubated at 56°C for 10 minutes. Ethanol was added and the lysate was added to a DNeasy Mini spin column where the DNA was bound to a membrane. The DNA was twice washed before being eluted. DNA concentrations were measured using a Nanodrop One (ThermoFisher).

### Immunocytochemistry (ICC)

Cells were fixed in 4% PFA in PBS for 10 minutes, washed in PBS, then incubated with blocking solution (10% goat serum, 2% bovine serum albumin, 0.2% triton in PBS) for 2 hours. Cells were then incubated in primary antibodies [MBP (1:50; Abcam; ab7349), OLIG2 (1:100; Atlas Antibodies; HPA003254), GFAP (1:500; Invitrogen; 14-9892-82), anti-Tubulin beta 3 (clone TUJ1; 1:1,000; BioLegend; 801202), IBA1 (1:500; Wako; 019-19741)] in block overnight at 4°C, washed in PBS, then incubated in secondary antibodies [Goat anti-Rat Alexa Fluor 647 (1:1,000; Invitrogen; A21247),Goat anti-Mouse Alexa Fluor 488 (1:1,000; Invitrogen; A11001), Goat anti-Rabbit Alexa Fluor 594 (1:1,000; Invitrogen; A11012), Goat anti-Rabbit Alexa Fluor 488 (1:1,000; Invitrogen; A11008)] for 1 hour at RT. Cells were then labelled with DAPI.

### qPCR

Total RNA was extracted from iPSC and iPSC-derived cortical neuron cells using a RNeasy Mini kit (QIAGEN; 74104), including a DNase treatment step, according to the manufacturer’s instructions. RNA was reverse-transcribed using a High-Capacity cDNA Reverse Transcription Kit (Applied Biosystems, Carlsbad, CA, USA) according to the manufacturer’s instructions. Quantitative analysis was carried out using qPCR with gene specific SYBR green forward (Fwd) and reverse (Rev) primers (*TBR1*: Fwd AGCAGCAAGATCAAAAGTGAGC, Rev ATCCACAGACCCCCTCACTAG; *CTIP2*: Fwd CTCCGAGCTCAGGAAAGTGTC, Rev TCATCTTTACCTGCAATGTTCTCC; *RPL18A*: Fwd CCCACAACATGTACCGGGAA, Rev TCTTGGAGTCGTGGAACTGC) using a Roche LightCycler 480 II. Gene expression in all samples was quantified using the ΔΔCt method normalised first to the housekeeping gene *RPL18A*, then to the average for all iPSC samples.

### ELISA

ELISAs for human Aβ40 and human Aβ42 were carried out using commercial kits (Aβ40: Invitrogen KHB3481; Aβ42: Invitrogen KHB3441) according to the manufacturer’s instructions. Briefly, samples were added to the wells with capture antibody prebound, together with detection antibody, and incubated for 3 hours. Wells were then washed before being incubated with anti-rabbit IgG HRP for 30 minutes. Wells were again washed, incubated with stabilized chromogen for 30 minutes, and the reaction stopped. Absorbance at 450 nm was measured using a FLUOstar Omega microplate reader.

### Immunodepletion of Aβ

Oligodendrocyte conditioned media was incubated with anti-Aβ antibody 6E10 (1:500; BioLegend; 803001) for 2 hours at 4°C. Samples were then incubated with prewashed Sepharose Protein A/G beads (50μl; Rockland; PAG50-00-0002) for 2 hours at 4°C. Beads with bound Aβ and antibody were then removed by centrifugation for 30 seconds at 12,500 RPM.

### Quantification of soluble Aβ aggregates

The technique used to quantify soluble aggregates has previously been validated in multiple studies and has been shown to specifically detect aggregates, and not monomers^17–19^. It relies on the binding of two copies of the same antibody (6E10, which only has a single epitope^20^), ensuring that what are detected are at least dimers. The clustering analysis used requires close proximity of at least two of these dimers, thus further ensuring that what are detected are aggregates. This method allows quantification of stable assemblies or aggregates at picomolar concentrations, which is a much higher sensitivity than bulk biochemical methods.

Single-molecule pull-down coverslip preparation was performed as previously described^21^. Briefly, neutravidin (0.2 mg/ml) in TBS with 0.05% Tween 20 (TBST) was added to glass coverslips covalently mounted with polyethylene glycol (PEG) and biotin for 10 minutes, followed by two wash steps with TBST and once with TBS with 1% Tween 20 (1%T). Afterwards, biotinylated 6E10 antibody (10nM; BioLegend; 803007) was added for 15 minutes, followed by two wash steps with TBST and once with 1%T. The media samples were added and incubated overnight at 4℃ followed by two wash steps with TBST and once with 1%T. The coverslips were then incubated with 6E10 antibody (500pM; BioLegend; 803020), labelled with single strand DNA^22^ (ACCACCA) for 45 minutes, followed by two wash steps with TBST and once with 1%T. After washing, TetraSpeck microspheres (1:7,000 in TBS, 10 µL, Thermo Scientific, Cat. T7279) were introduced to each well for 10 minutes. The TetraSpeck solution was then removed, followed by 2x wash with TBST, and a second PDMS gasket (Merck, GBL-103250-10EA) was stacked on the coverslip before introducing 3 µL of complimentary imaging strand (TGGTGGT-cy3B; atdbio) in TBS. Finally, the coverslip was sealed with another coverslip on top of the second PDMS gasket. Then the coverslips were imaged on a purpose built^21^ TIRF microscope using a 520 nm laser, with 100 milliseconds of exposure for 4,000 frames for each field of view. 3 fields of view were acquired from each well. Acquired frames were stacked, reconstructed, drift corrected, and analysed for the number of aggregates in each field of view, as well as aggregate length using the ACT software^19^. Images of individual aggregates were generated using the ASAP software^23^.

### Analysis of single nucleus RNA sequencing data

Human single nucleus RNA sequencing data from Zhou *et al.*, 2020^24^ were obtained via the AD Knowledge Portal (https://adknowledgeportal.org) study snRNAseqAD_TREM2. Human single nucleus RNA sequencing data from Bakken *et al.*, 2021^25^ were obtained from the Human Protein Atlas (proteinatlas.org). Human single nucleus RNA sequencing data from Mathys *et al.*, 2019^26^ and Lake *et al.*, 2018^27^ were obtained from The Myeloid Landscape 2 portal (http://research-pub.gene.com/BrainMyeloidLandscape)^28^. Analysis of Zhou *et al.*, 2020 dataset: Data analysis was performed using R (4.1.2) and packages Seurat (4.0.6), dplyr (1.0.7), ggplot2 (3.3.5), pheatmap (1.0.12), and wesanderson (0.3.6). For analysis, we retained all cells included in the labelled metadata accompanying the study, without further quality filter. We performed data normalization, variable gene identification, data scaling, dimensionality reduction with PCA and clustering analysis using functions implemented in the package Seurat. Subgroups Ex0 and Ex1 were considered together for excitatory neurons, and subgroups Oli0 and Oli1 were considered together for oligodendrocytes. The z-score was calculated from the log_2_ (normalised counts + 1) and plotted as a heat map for each dataset. Analysis of other 3 datasets: Data for each gene of interest were obtained as normalised counts averaged for each cell type. The z-score was calculated from the log_2_ (normalised counts + 1) and plotted as a heat map for each dataset.

### Analysis of proteomics data

Protein expression data of isolated mouse brain cells was generated by Sharma *et al.,* 2015^29^ using Magnetic-Activated Cell Sorting (MACS) to isolate microglia, astrocytes, oligodendrocytes and neurons from the brains of young (P8) C57/BL6 mice (3 biological replicates), analysed by mass spectrometry. These data were obtained from the journal website. Data for each protein of interest were obtained as log_2_ LFQ intensity averaged for each cell type. The z-score was calculated from the log_2_ LFQ intensity and plotted as a heat map.

### Imaging and analysis

IHC: for oligodendrocyte analysis, sections were imaged using a Zeiss LSM 800 confocal microscope using a 40x magnification (field of view (FOV): 159.73 x 159.73 μm; 2101 x 2101 pixels), 4-6 FOV from Layers 5/6 of the cortex were captured across 2-3 sections, and cells were quantified manually using the FIJI build of ImageJ (open source) after blinding the experimenter to animal genotype. For APP and BACE analysis, sections were imaged using a Zeiss LSM 980 confocal microscope using a 40x magnification, 3-4 FOV from the cortex were captured, and cells were quantified manually using the FIJI build of ImageJ (open source). For plaque analysis, sections were imaged using a Zeiss AxioScan slide scanner, capturing the whole of 3 sagittal sections per animal, with a maximum resolution of 0.325 μm/pixel. Images were first manually segmented to select regions of interest [all visible cortical areas (comprising of visual, retrosplenial and M2 motor areas), Layers 2/3 of motor cortex (M2), Layers 5/6 of motor cortex (M2), Layers 2/3 of retrosplenial cortex, Layers 5/6 of retrosplenial cortex, corpus callosum (including splenium), area CA1 of hippocampus] and remove regions with post-mortem tissue damage or poor imaging quality. The boundary between Layers 2/3 and Layers 5/6 was identified based on the sharp reduction in the density of cell nuclei. Analysis was then automated using the FIJI build of ImageJ software (open source) to segment, count and measure Aβ^+^ plaques. Myelin sheath lengths were analysed as previously described^30,31^: images were captured using a Leica TCS SPE confocal microscope at 20x magnification with a z-step size of 2μm. 4 images were collected from Layer 2/3 of the primary somatosensory cortex from each of 3 sections per animal. Myelin sheaths were measured using the simple neurite trace plug-in for the FIJI build of ImageJ (open source). Only complete myelin sheaths that started and ended within the 30μm section were traced and measured. 120 sheaths were measured per mouse. Imaging and analysis was performed blind to genotype/condition. Oligodendrocyte autophagy was analysed as previously described^32,33^: images were captured using a Zeiss LSM 980 confocal microscope using a 63x magnification (pixel size 0.132μm) with a z-step size of 0.5μm, 4 FOV were captured per animal from Layers 5/6 of the cortex, with 2-3 cells analysed per FOV. Cells were segmented based on Olig2 staining, the nuclear mask was dilated to capture the cytoplasm, and the fluorescent intensity normalised to surface was measured. Imaging and analysis was performed blind to genotype/condition.

RNAscope: sections were imaged using a Zeiss LSM 880 or Zeiss LSM 980 confocal microscope using a 40x magnification (field of view (FOV): 212.55 x 212.55 μm; 1024 x 1024 pixels). 10 FOV were captured from each brain, 5 corresponding to Layer 5/6 of the cortex from different parts of the tissue section and 5 corresponding to Layer 2/3. The number of cells expressing different genes was quantified manually using the FIJI build of ImageJ software (open source) after blinding of the experimenter to the donor ID and diagnosis. Densitometry was carried out using a custom macro in the FIJI build of ImageJ software to segment cell nuclei (rather than algorithmically inferring cell boundaries, which is liable to introduce bias when comparing cell types which greatly differ in size), classify them as *MBP*^+^ or *RBFOX3*^+^, and then measure the number and area of BACE1 and APP spots in each cell.

ICC: cells were imaged using a Zeiss Cell Discoverer 7 imaging system with an LSM 900 confocal imaging module using a 20x magnification (field of view (FOV): 202.45 x 202.45 μm; 1024 x 1024 pixels). For each line, a minimum of 3 FOV were captured from each of 2 wells. Cell number was quantified manually using a custom macro in the FIJI build of ImageJ software, while cell positivity was assessed manually.

### Statistics

Graphs were plotted and statistical tests were carried out using GraphPad Prism software or Matlab. Statistical tests were chosen for each experiment based on the normality of data and sample matching. Actual statistical tests used for each experiment are stated in the figure legends. All statistical tests used were two-sided unless otherwise stated. n values stated refer to true biological replicates – depending on the experiment an individual n is a human brain (comprising 5 FOV imaged), a cell line (comprising 1-4 independent inductions, each of which is the average of 2-5 wells), a mouse (comprising 2-3 whole sagittal sections imaged or 3-12 FOV), or a neuron/unit. Actual n values and what they refer to for each experiment are stated in the figure legends.

