## Supplementary Information for "Selective suppression of oligodendrocyte-derived amyloid beta rescues neuronal dysfunction in Alzheimer’s Disease"

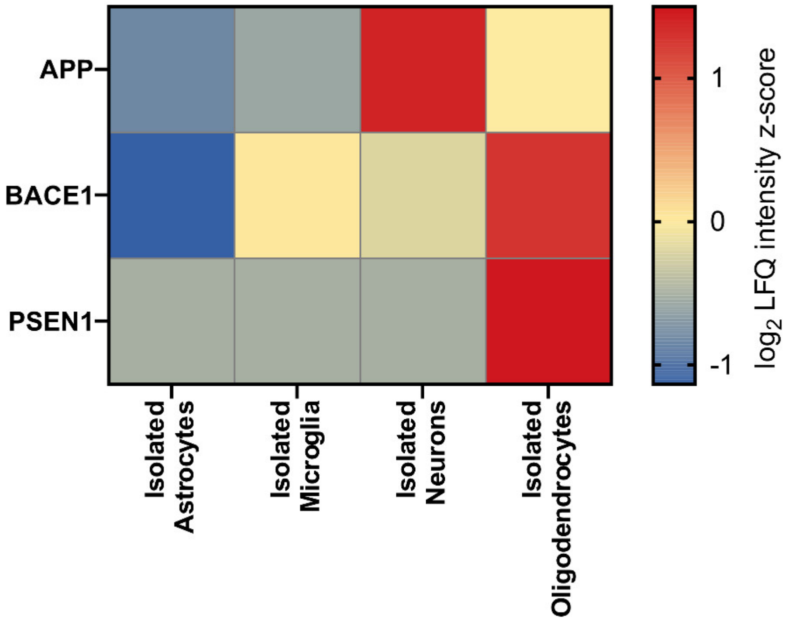


**Supplementary Fig. 1: Proteins required to produce Aβ are found at high levels in oligodendrocytes.** Heatmap showing the $\log_2$ (LFQ intensity) z-score of proteins of interest from isolated mouse astrocytes, microglia, neurons and oligodendrocytes shows high amounts of APP, BACE1 and PSEN1 in isolated oligodendrocytes. Proteomics data from Sharma *et al*., 2015[1] was generated from cells isolated by Magnetic-Activated Cell Sorting (MACS) from C57/BL6 mice.

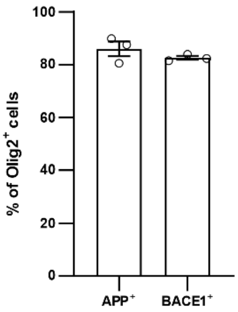


**Supplementary Fig. 2: Proteins required to produce Aβ are found in the majority of oligodendrocytes.** Quantification of immunofluorescent images from 4-month-old wild type mice (see **Fig. 1e-f**) showing the percentage of Olig2$^+$ cells which are APP$^+$ or BACE1$^+$. Each data point represents an individual mouse (n=3) with bars showing mean ± SEM.

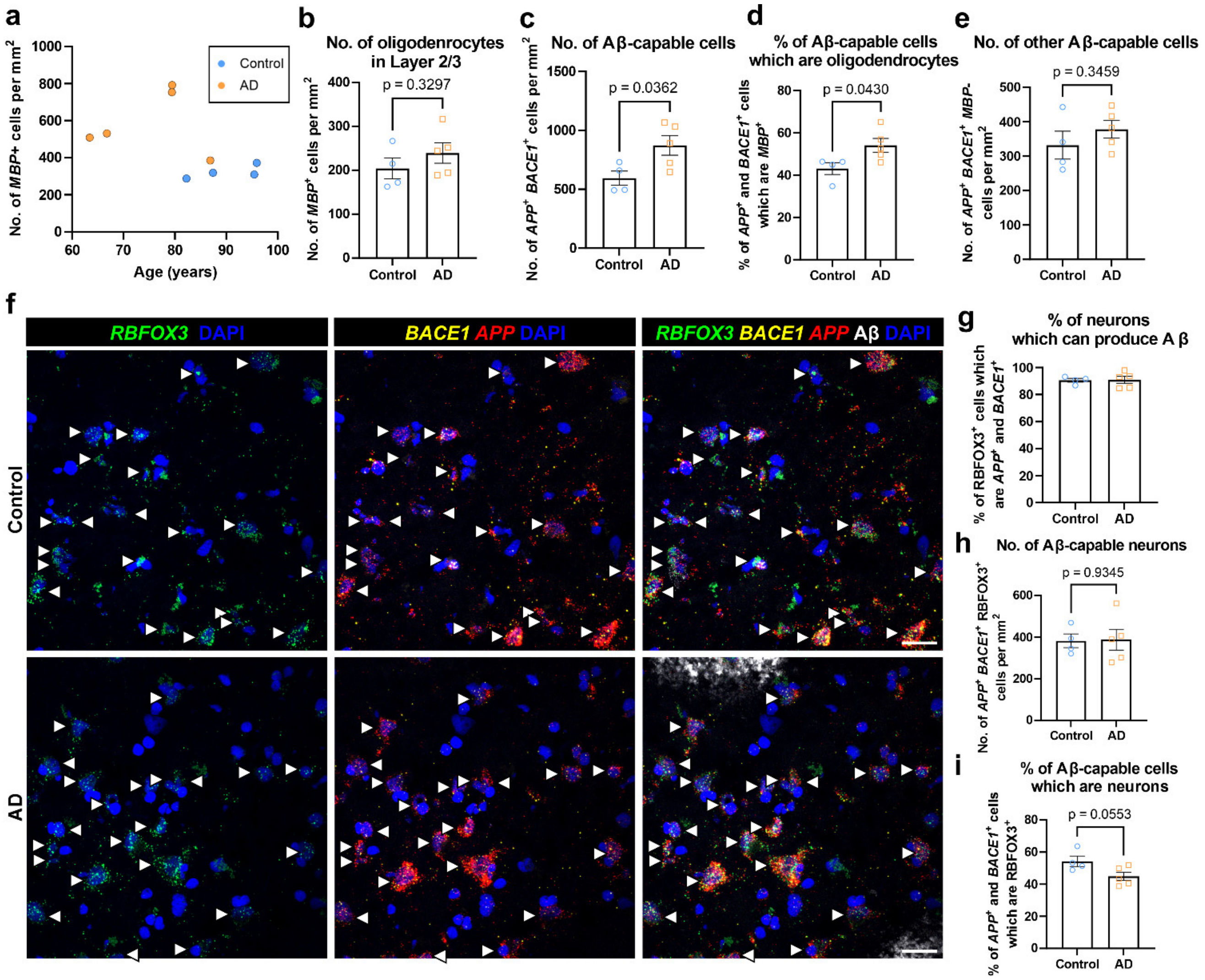


**Supplementary Fig. 3: Increase in cells capable of producing Aβ in brains from patients with sporadic AD brains is largely due to oligodendrocytes. a,** Plot showing the number of *MBP*$^+$ cells found in Layers 5/6 (**Fig. 1a, c**) of the prefrontal cortex against the age of the donor. A linear regression analysis indicated no significant relationship ($F(1,7)$=1.958, $p$ = 0.2045), with a low coefficient of determination ($r^2$=0.2185), suggesting that age is not a significant contributor to the number of *MBP*$^+$ cells in Layers 5/6 of the prefrontal cortex. **b,** Quantification showing no significant change in the number of oligodendrocytes in Layer 2/3 of the prefrontal cortex of AD brains. **c,** Quantification showing a significant increase in the number of all Aβ producing cells in sporadic AD (sAD) brains compared to controls. **d,** Quantification showing a significant increase in the proportion of Aβ producing cells which are oligodendrocytes in sAD brains. **e,** Quantification of the number of Aβ producing cells which are not oligodendrocytes showing no significant difference between control and AD brains. **f,** Fluorescent images from Layers 5/6 of control (top) and sporadic AD (sAD; bottom) post-mortem human prefrontal cortex labelled for *RBFOX3* (neuron-specific gene; green), *BACE1* (yellow), *APP* (red), Aβ (identified by 6E10-antibody; white), and DAPI (nuclei; blue). Aβ-capable neurons (*RBFOX3*$^+$ *BACE1*$^+$ *APP*$^+$ nuclei) are marked with white arrowheads. Note the high variability in expression levels of *APP* and *BACE1* between neurons, with high expression in some cells but minimal expression in others. Scale bar = 25µm. **g,** Quantification of the proportion of neurons which are capable of producing Aβ. **h,** Quantification of the number of neurons which are capable of producing Aβ. **i,** Quantification showing the proportion of Aβ-capable cells which are neurons. Each data point represents a single brain (n=4 control brains, n=5 sAD brains) with bars representing mean ± SEM. Unpaired t-test; $t(7)$=1.047, 2.585, 2.467, 1.011, 0.08304, 0.08520, 2.297 in **b,c,d,e,g,h,i** respectively.

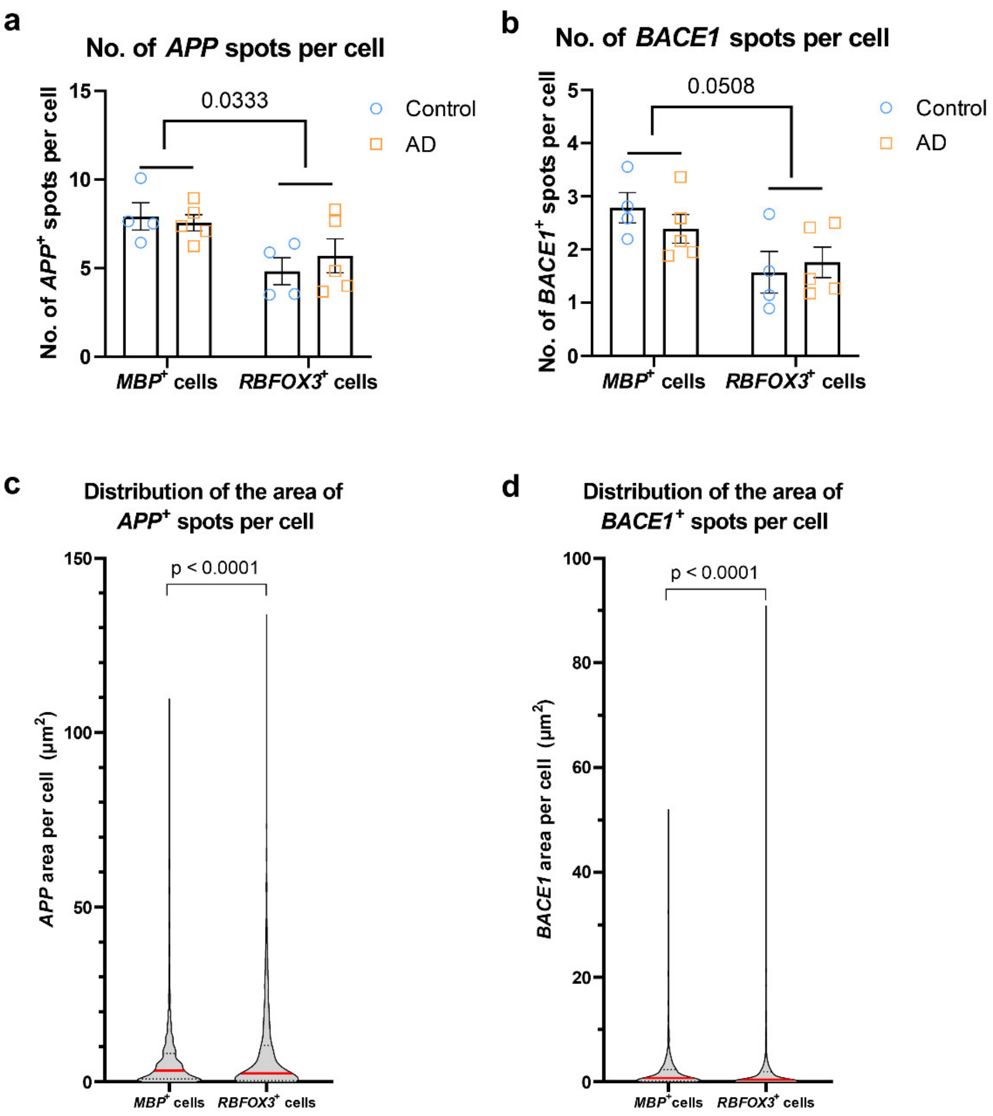


**Supplementary Fig. 4: Densitometric analysis shows increased *APP* and *BACE1* in oligodendrocytes compared to neurons. a-b,** Quantification showing more $APP^+$ (**a**) and $BACE1^+$ (**b**) spots per cell in oligodendrocytes ($MBP^+$ cells) compared to neurons ($RBFOX3^+$ cells) in both control and sAD human post-mortem brains. Each data point represents a single brain (n=4 control brains, n=5 sAD brains) with bars representing mean ± SEM. Two-way repeated measures ANOVA. Cell type effect: **a**: $F(1,7)=6.979$; **b**: $F(1,7)=5.540$. **c-d,** Violin plots demonstrating the greater variability in expression levels of *APP* (**c**) and *BACE*1 (**d**) in neurons ($RBFOX3^+$ cells) compared to oligodendrocytes ($MBP^+$ cells) in human post-mortem brains (n=843 $MBP^+$ cells, n=1100 $RBFOX3^+$ cells from 4 control and 5 sAD brains). Red lines indicate mean, while dotted black lines indicate quartiles. Fligner-Killeen test for equality of variances: $FK(1,1944) = 388.69, 533.44$ in **c, d** respectively.

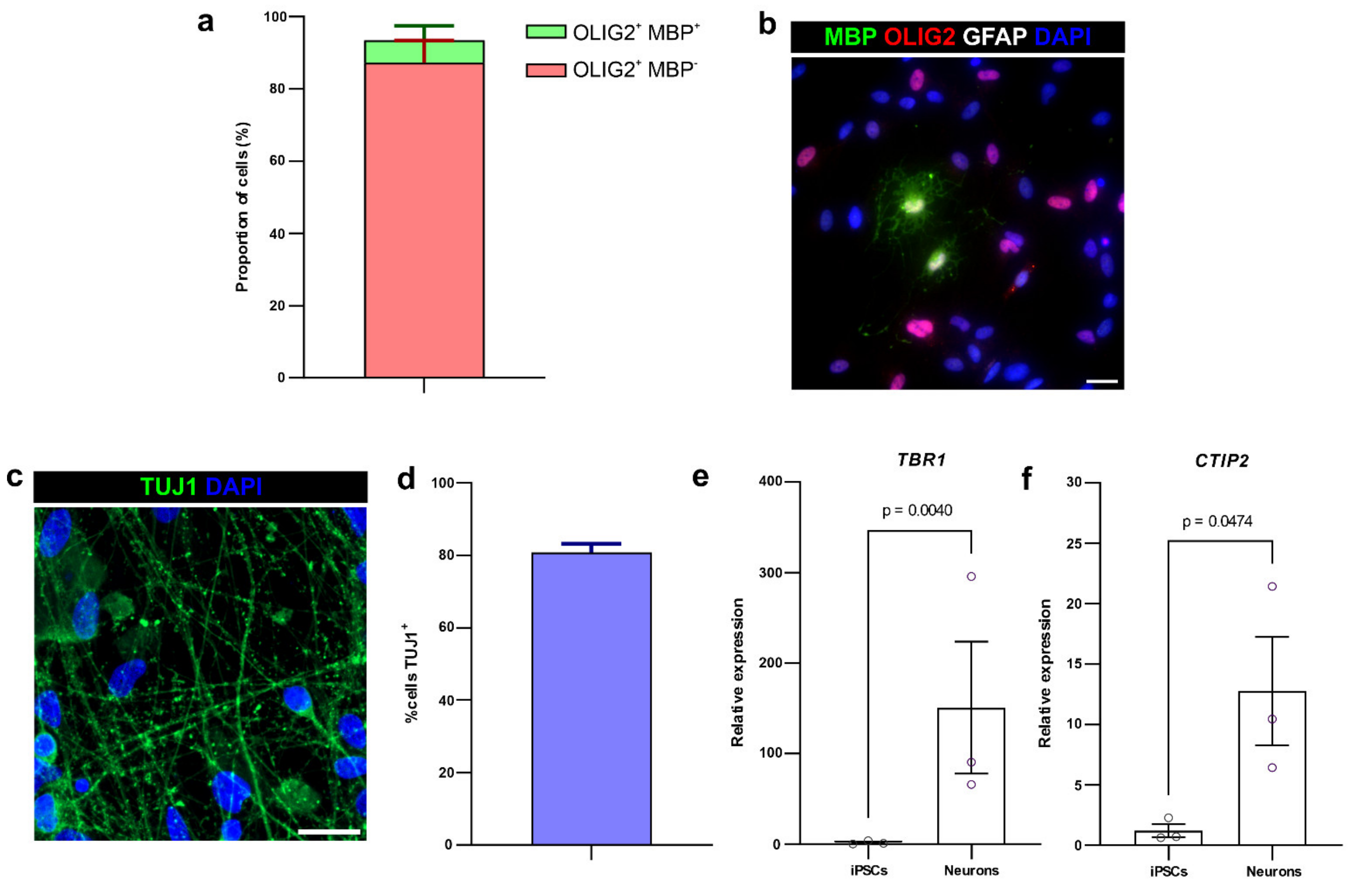


**Supplementary Fig. 5: Human iPSC-derived oligodendrocyte and neuron cultures have high purity. a,** Quantification of the proportion of cells in oligodendrocyte cultures which are oligodendrocyte precursor cells (OPCs; $OLIG2^+$ $MBP^-$) and mature, myelin-expressing oligodendrocytes ($OLIG2^+$ $MBP^+$) shows that 93.5% ± 2.2% of all cells in these cultures are either oligodendrocytes or OPCs. Bars show mean + SEM. n=3 cell lines (one induction per line). **b,** Representative fluorescent image of human iPSC-derived oligodendrocyte culture immunolabelled for MBP (green), OLIG2 (red), GFAP (white) and DAPI (blue). No $GFAP^+$ astrocytes were found in 3 out of 3 cultures examined. Scale bar = 25µm. **c,** Fluorescent image of human iPSC-derived neuronal culture immunolabelled for the neuronal marker TUJ1 (green), and DAPI (nuclei; blue). Scale bar = 25µm. **d,** Quantification of the proportion of cells in neuronal cultures which are neurons shows a high proportion of $TUJ1^+$ cells, consistent with previous studies [2]. Bar shows mean + SEM. n=3 cell lines (one induction per line). **e-f**, qPCR data showing high expression of deep cortical layer markers *TBR1* (**e**) and *CTIP2* (**f**) in neuronal cultures compared to undifferentiated iPSCs. Graphs show relative expression of the gene of interest, normalised to the average for iPSC cultures using the ΔΔCt method (with *RPL18A* as the housekeeping gene). Each data point represents a different cell line (n=3; one induction per line) with bars showing mean ± SEM. Ratio paired t-test: $t(2)=15.74$, 4.430 in **e,f** respectively.

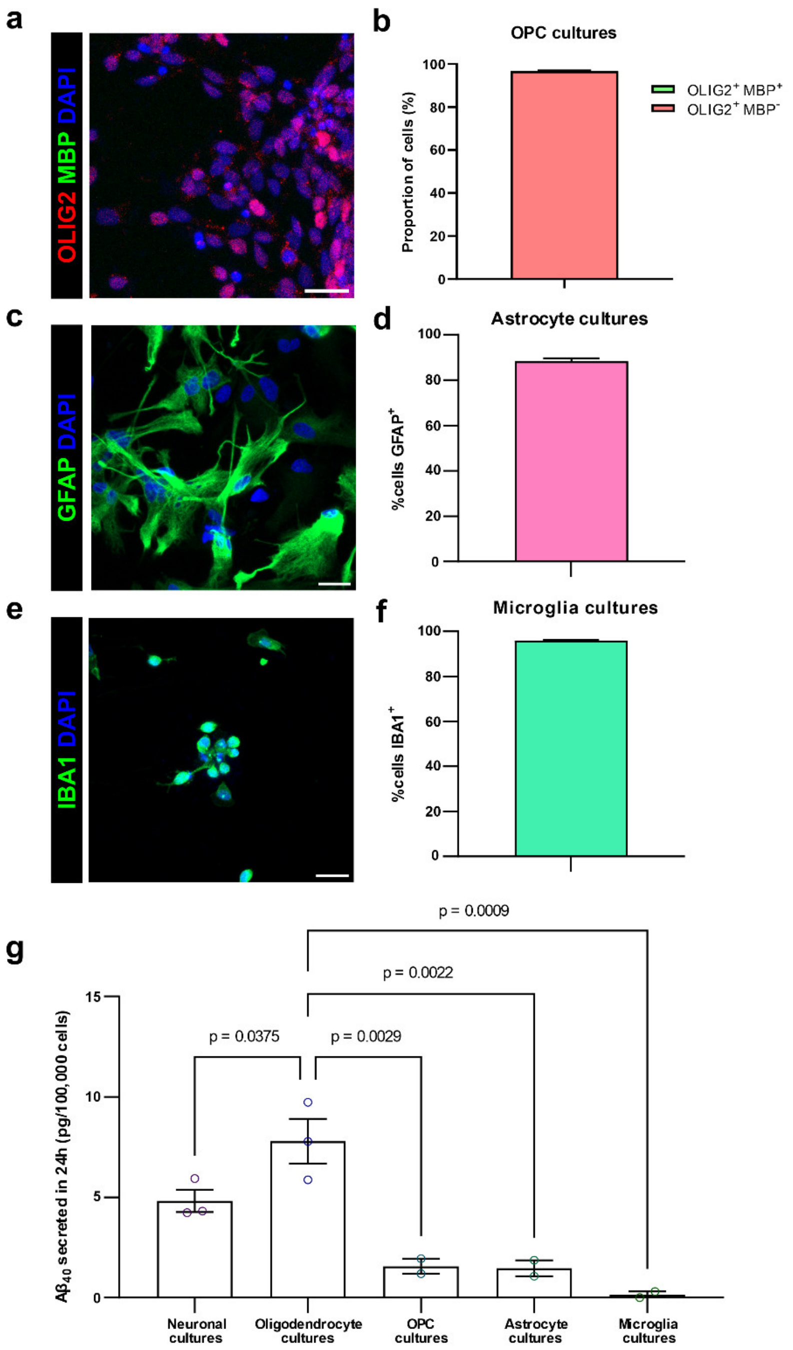


**Supplementary Fig. 6: Immature OPCS, astrocytes and microglia produce low amounts of Aβ. a-f,** Representative fluorescent images (**a, c, e**) and quantifications showing characterisation of OPC (**a-b**), astrocyte (**c-d**) and microglia (**e-f**) cultures. In **a,** cells are immunolabelled for MBP (myelin basic protein; green), OLIG2 (marker of all oligodendroglia; red), and DAPI (nuclei; blue), with quantification in **b** showing 96.8% ± 0.2% of cells are $OLIG2^+$ $MBP^-$ OPCs while there are no mature $MBP^+$ oligodendrocytes. In **c,** cells are immunolabelled for GFAP (green) and DAPI (blue), with quantification in **d** showing 88.3% ±1.0% of cells are $GFAP^+$ astrocytes. In **e,** cells are immunolabelled for IBA1 (green) and DAPI (blue), with quantification in **f** showing 95.9% ± 0.2% of cells are $IBA1^+$ microglia. In **a,c,e,** scale bar = 25µm. In **b,d,f,** bars show mean + SEM, n=2 cell lines (one induction per line). **g,** ELISA data showing very low amounts of $A\beta_{40}$ produced by human iPSC-derived OPCs, astrocytes and microglia compared to neurons and oligodendrocytes. Each data point represents the average of two (OPCs, astrocytes, microglia) or three (neurons, oligodendrocytes) independent inductions/harvests from a different cell line (OPCs, astrocytes, microglia: n=2; neurons, oligodendrocytes: n=3), with bars representing mean ± SEM. Mixed effects analysis ($F(4,5)=27.77$, $p=0.013$) with Dunnet's post-hoc tests.

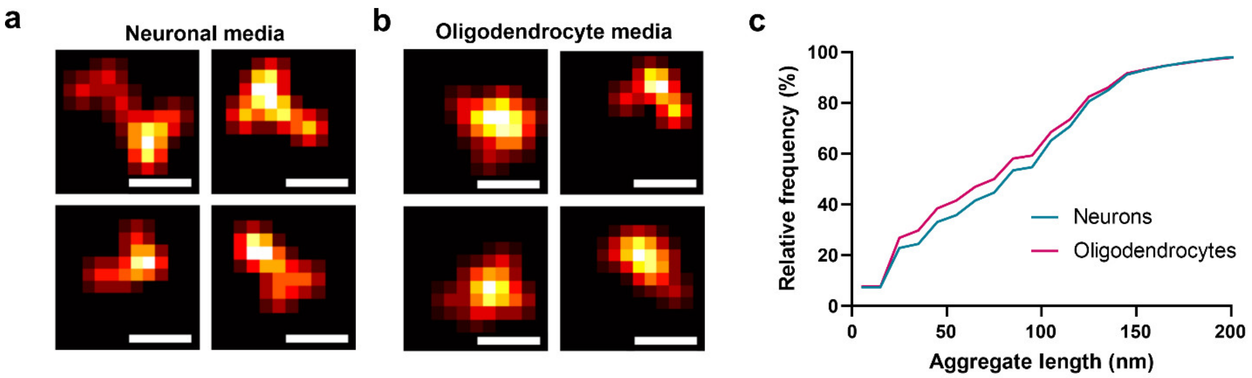


**Supplementary Fig. 7: A majority of soluble Aβ aggregates produced by human neurons and oligodendrocytes are between 20nm and 200nm. a,b,** Examples of super-resolved aggregates detected in neuronal media (**a**) and oligodendrocyte media (**b**). Scale bar = 50nm. **c,** Cumulative frequency histogram showing the size distribution of aggregates produced by neurons and oligodendrocytes (bin size = 10nm).

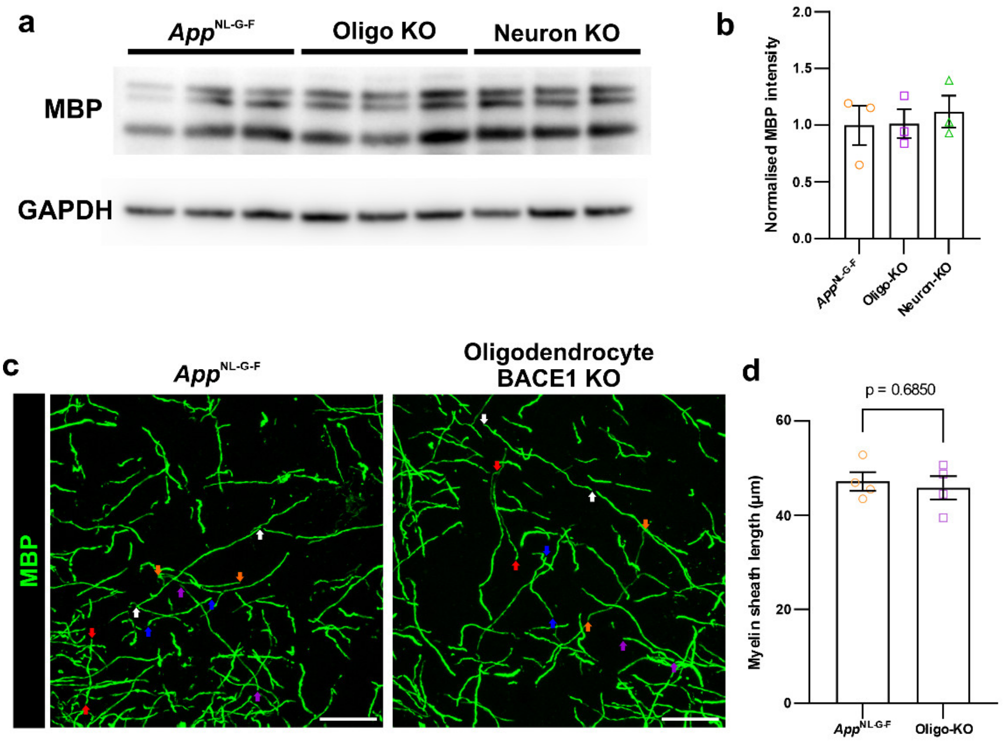


**Supplementary Fig. 8: Genetic suppression of BACE1 in oligodendrocytes after 4 weeks of age does not affect total amount of MBP or myelin sheath length. a,** Western blot for MBP (top) and GAPDH loading control (bottom) on forebrain homogenates from *App*$^{\text{NL-G-F}}$ mice (left 3 lanes), and *App*$^{\text{NL-G-F}}$ mice with BACE1 knocked out (KO) specifically in oligodendrocytes (Oligo-KO; middle 3 lanes) or neurons (Neuron-KO; right 3 lanes). **b,** Quantification of the MBP bands normalised to GAPDH loading control shows no significant difference in MBP levels in mice with BACE1 knocked out. In **b,** data points represent individual mice (n=3 per group) with bars showing mean ± SEM. One-way ANOVA: $F(2,6)=0.1924$, $p=0.8299$. **c,** Representative immunofluorescent images of myelin sheaths (labelled with MBP, green) in the sparsely myelinated Layers 2/3 of the primary somatosensory cortex showing no significant differences between *App*$^{\text{NL-G-F}}$ mice (left) and *App*$^{\text{NL-G-F}}$ mice with BACE1 KO specifically in oligodendrocytes (Oligo-KO; right). Coloured pairs of arrows indicate the start and end of example myelin sheaths. Scale bar = 30µm. **d,** Quantification of the lengths of myelin sheaths in the cortex, as previously described[3,4], shows no differences in mice with BACE1 knocked out after 4 weeks of age. In **d,** each data point represents the average of 120 myelin sheaths measured from an individual mouse (n=4 per group) with bars showing mean ± SEM. Unpaired t-test: $t(6)=0.4259$.

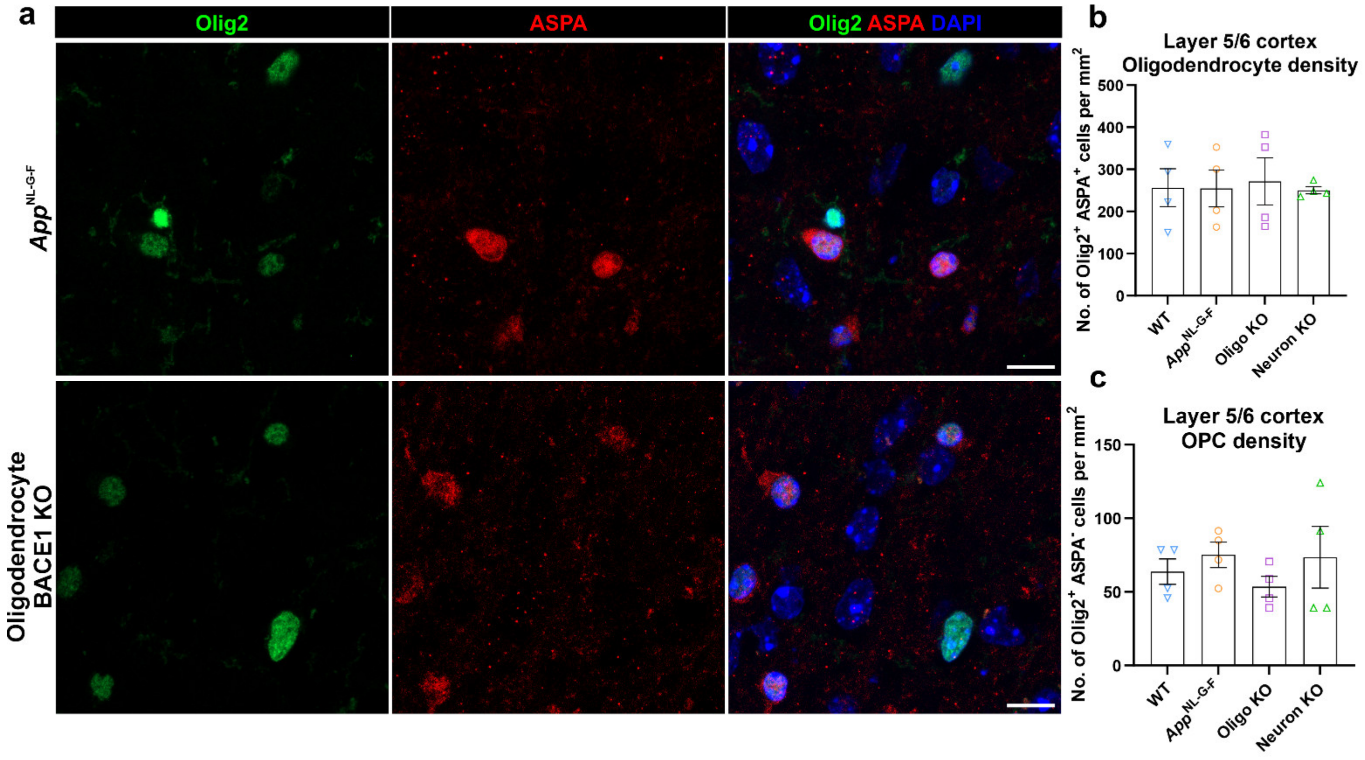


**Supplementary Fig. 9: Genetic suppression of BACE1 in oligodendrocytes after 4 weeks of age has no significant effect on oligodendrocyte number. a,** Immunofluorescent images showing pan-oligodendroglial marker Olig2 (green), oligodendrocyte-specific marker ASPA (red) and DAPI (blue) in the cortex of *App*$^{NL\text{-}G\text{-}F}$ control mice (top) and *App*$^{NL\text{-}G\text{-}F}$ mice with BACE1 KO specifically in oligodendrocytes (bottom). Scale bar = 25µm. **b,** Quantification of Olig2$^{+}$ ASPA$^{+}$ cells shows no significant difference in oligodendrocyte number in mice with BACE1 KO. **c,** Quantification of Olig2$^{+}$ ASPA$^{-}$ cells shows no significant difference in the number of oligodendrocyte precursor cells (OPCs) in mice with BACE1 KO. In **b,c,** data points represent individual mice (n=4 per group) with bars showing mean ± SEM. One-way ANOVA: $F$(3,12)=0.04745, 0.6279; $p$=0.9856, 0.6107 in **b,c** respectively.

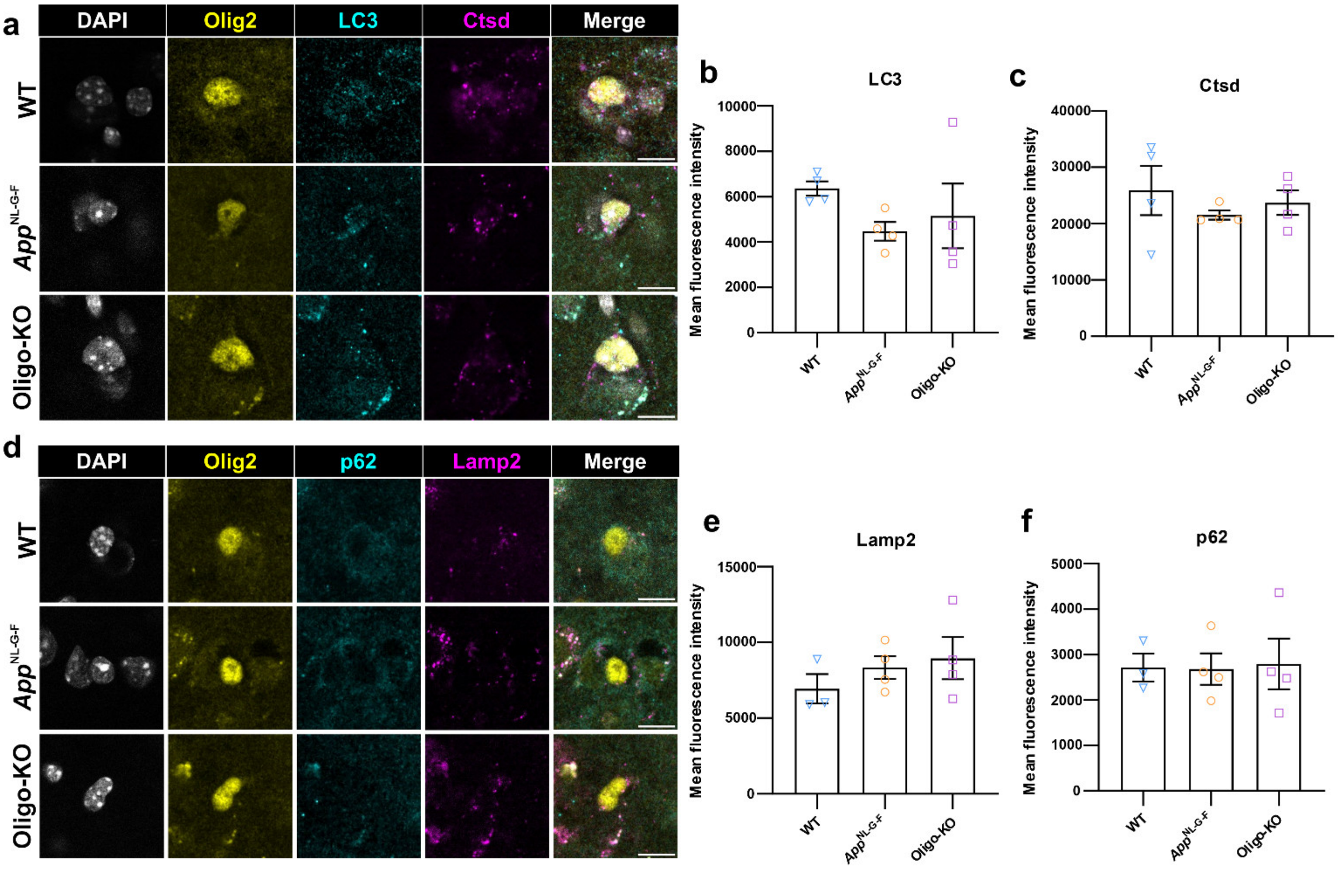


**Supplementary Fig. 10: Genetic suppression of BACE1 in oligodendrocytes of *App*$^{\text{NL-G-F}}$ mice after 4 weeks of age has no significant effect on oligodendrocyte autophagy[5,6]. a,** Immunofluorescent images showing oligodendroglial marker Olig2 (yellow), autophagosome marker MAP1LC3B/LC3B (microtubule-associated protein 1 light chain β; LC3; cyan), lysosome protease Cathepsin-d (Ctsd; magenta) and nuclear marker DAPI (white) in the cortex of WT (top), *App*$^{\text{NL-G-F}}$ control mice (middle) and *App*$^{\text{NL-G-F}}$ mice with BACE1 KO specifically in oligodendrocytes (Oligo-KO; bottom). **b-c,** Quantification shows no significant changes in LC3 (**b**), or Ctsd (**c**) within oligodendroglia upon knockout of BACE1 in oligodendrocytes. **d,** Immunofluorescent images showing oligodendroglial marker Olig2 (yellow), autophagy adapter Sqstm1/p62 (cyan), lysosome marker Lamp2 (lysosome-associated membrane protein 2; magenta) and nuclear marker DAPI (white) in the cortex of WT (top), *App*$^{\text{NL-G-F}}$ control mice (middle) and *App*$^{\text{NL-G-F}}$ mice with BACE1 KO specifically in oligodendrocytes (Oligo-KO; bottom). **e-f,** Quantification shows no significant changes in p62 (**e**), or Lamp2 (**f**) within oligodendroglia upon knockout of BACE1 in oligodendrocytes. In **b-c, e-f,** data points represent individual mice (n=3 WT in **e-f**, n=4 for all other groups) with bars showing mean ± SEM. One-way ANOVA: $F(2,9)=1.207, 0.5826$; $p=0.3434, 0.5782$ in **b,c** respectively. $F(2,6)=0.7969, 0.01823$; $p=0.4835, 0.9820$ in **e,f** respectively.

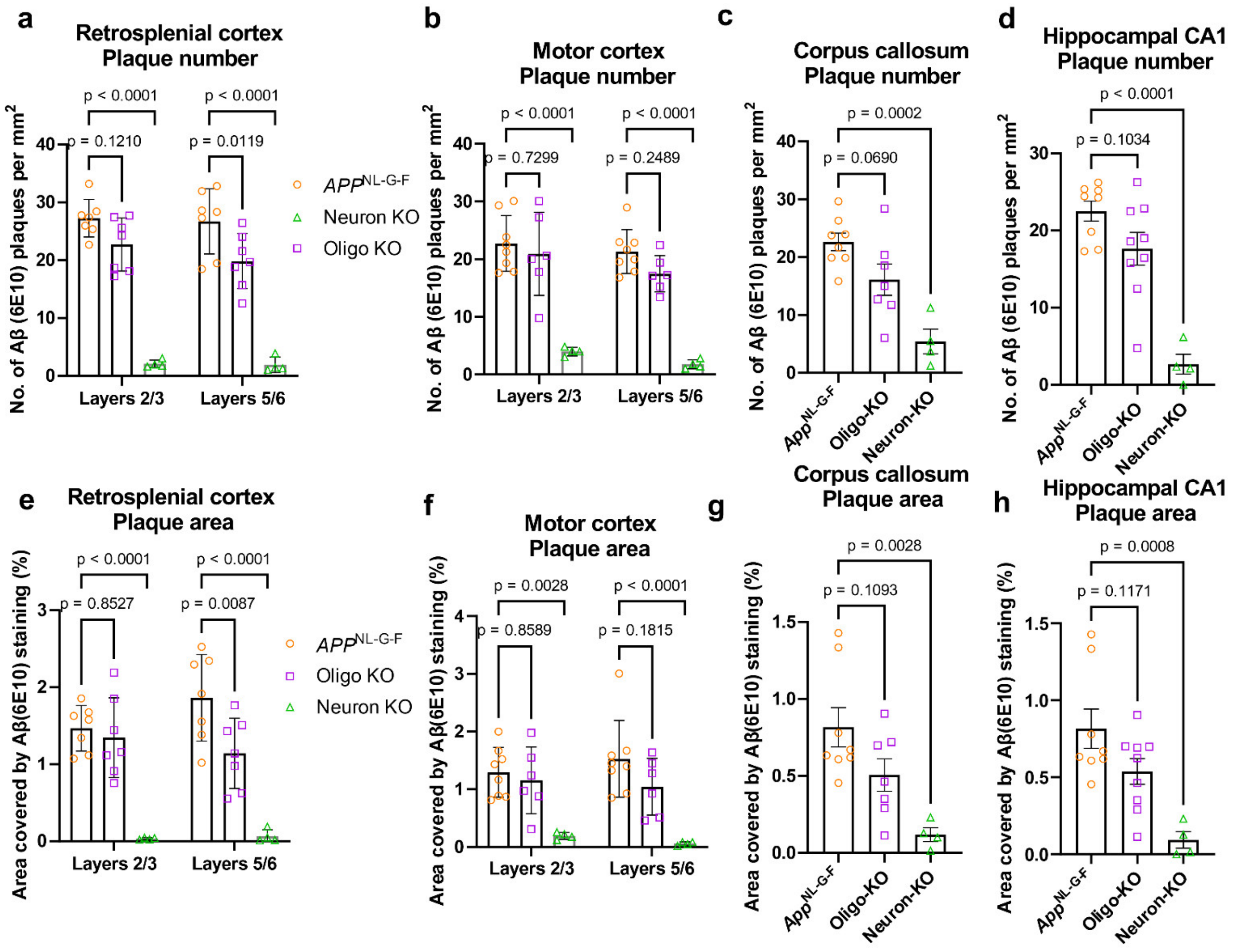


**Supplementary Fig. 11: Genetic suppression of oligodendrocyte Aβ production leads to a modest reduction in Aβ plaque burden while genetic suppression of neuronal Aβ production eliminates plaques in all regions examined of the *App*$^{\text{NL-G-F}}$ mouse model of AD. a,e** Quantification of plaque number (**a**) and plaque area (**e**) across different layers of the retrosplenial cortex suggests that oligodendrocyte specific KO of BACE1 leads to a greater reduction in plaque pathology in deeper layers (Layers 5/6) than is observed in superficial layers (Layers 2/3). Data points represent individual mice (n=7 *App*$^{\text{NL-G-F}}$, 7 Oligo-KO, 4 Neuron-KO) with bars showing mean ± SEM. Two-way repeated measures ANOVA with Tukey's post-hoc tests. BACE1 KO effect: $F(2,15)=61.25$ (**a**), 22.59 (**e**); $p<0.0001$ (all); Layer-BACE1 KO interaction effect: $F(2,15)=0.8174$ (**a**), 5.892 (**e**); $p=0.4603$ (**a**), $p=0.0129$ (**e**). **b,f,** Quantification of plaque number (**b**) and plaque area (**f**) across different layers of the motor cortex. Data points represent individual mice (n=8 *App*$^{\text{NL-G-F}}$, 6 Oligo-KO, 4 Neuron-KO) with bars showing mean ± SEM. Two-way repeated measures ANOVA with Tukey's post-hoc tests. BACE1 KO effect: $F(2,15)=47.58$ (**b**), 13.24 (**f**); $p<0.0001$ (**b**), $p=0.0005$ (**f**); Layer-BACE1 KO interaction effect: $F(2,15)=0.2164$ (**b**), 0.9289 (**f**); $p=0.8079$ (**b**), $p= 0.4166$ (**f**). **c,d,g,h,** Quantification of plaque number (**c,d**) and plaque area (**g,h**) in the corpus callosum (**c,g**) and hippocampal area CA1 (**d,h**) shows a trend towards reduction in Oligo-KO mice in these regions. In **c,g,** data points represent individual mice (n=8 *App*$^{\text{NL-G-F}}$, 7 Oligo-KO, 4 Neuron-KO) with bars showing mean ± SEM. One-way ANOVA with Dunnet's post-hoc tests: $F(2,16)=12.94$, 7.549; $p=0.0005$, 0.0049 in **c,g** respectively. In **d,h,** data points represent individual mice (n=8 *App*$^{\text{NL-G-F}}$, 9 Oligo-KO, 4 Neuron-KO) with bars showing mean ± SEM. One-way ANOVA with Dunnet's post-hoc tests: $F(2,18)=21.83$, 9.338; $p<0.0001$, $p=0.0017$ in **g,h** respectively.

a

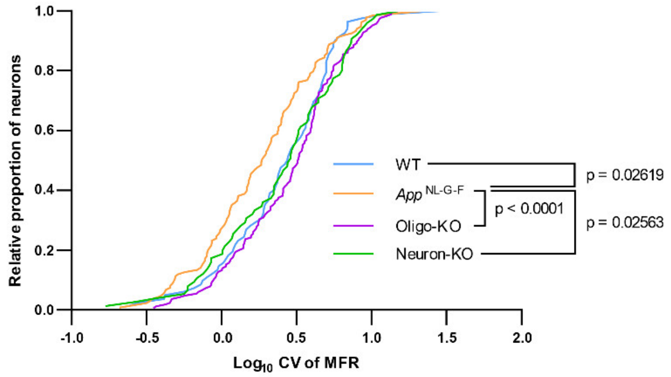


b

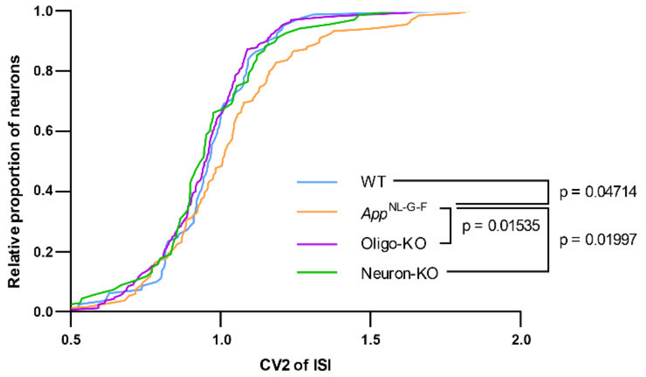


**Supplementary Fig. 12: Oligodendrocyte-specific knockout of BACE1 in *App*$^{NL-G-F}$ mice restores the temporal structure of neuronal firing to levels seen in WT controls. a,** Cumulative frequency distribution of the coefficient of variation (CV) of the mean firing rate (MFR) shows that variability in MFR is restored to WT levels in *App*$^{NL-G-F}$ mice with BACE1 knocked out (KO) specifically in oligodendrocytes (Oligo-KO; WT vs. Oligo-KO: $p = 0.6192$). **b,** Cumulative frequency distribution of the coefficient of variation (CV2) of the inter-spike interval (ISI) shows the restoration of spike train variability to WT levels in Oligo-KO mice (WT vs. Oligo-KO: $p = 0.5032$). In **a,** individual neurons/units are plotted (n=82 WT, 134 *App*$^{NL-G-F}$, 165 Oligo-KO, 75 Neuron-KO) from 4 (WT) or 3 mice per group. Kolmogrov-Smirnov tests ($D$=0.2026 (WT vs. *App*$^{NL-G-F}$), 0.2688 (*App*$^{NL-G-F}$ vs. Oligo-KO), 0.2088 (*App*$^{NL-G-F}$ vs. Neuron-KO)). In **b,** individual neurons/units are plotted (n=81 WT, 134 *App*$^{NL-G-F}$, 165 Oligo-KO, 68 Neuron-KO) from 4 (WT) or 3 mice per group. Kolmogrov-Smirnov tests ($D$=0.1891 (WT vs. *App*$^{NL-G-F}$), 0.1787 (*App*$^{NL-G-F}$ vs. Oligo-KO), 0.2215 (*App*$^{NL-G-F}$ vs. Neuron-KO)).

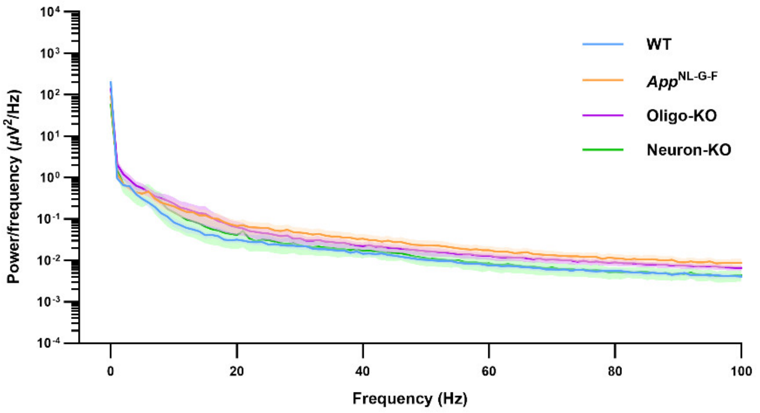


**Supplementary Fig. 13: No significant effect on local field potentials following BACE1 knockout in *App*$^{NL\text{-}G\text{-}F}$ mice.** Power spectrum of local field potential (LFP) recordings in retrosplenial cortex showing no major differences between genotypes. n=4 (WT), 3 (other groups) mice. One-way ANOVA: $F(3,9)=0.6235$, $p=0.6175$.

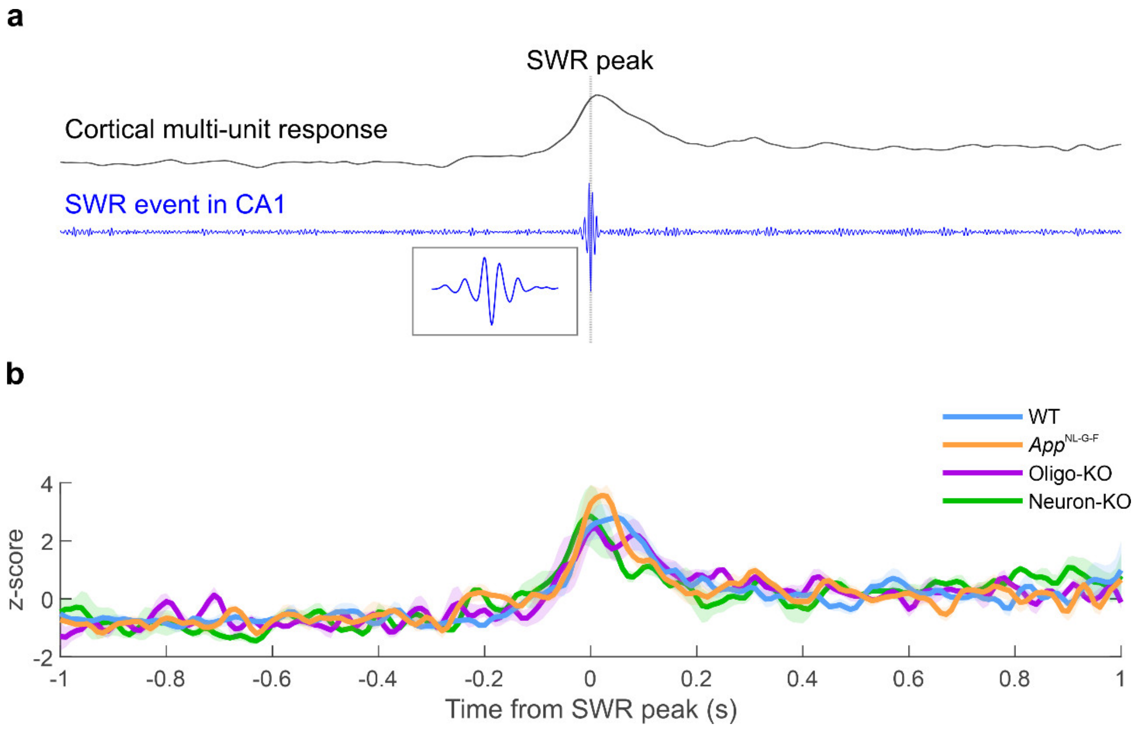


**Supplementary Fig. 14: No significant effect on cortical responses to sharp-wave ripple events in CA1 following BACE1 knockout in *App*$^{NL\text{-}G\text{-}F}$ mice. a,** Schematic illustrating a typical cortical multi-unit response (top; black) to a sharp wave ripple (SWR) event in CA1 (bottom; blue). **b,** Trace of the averaged cortical multi-unit responses to CA1 SWR events showing no significant differences between genotypes in amplitude [peak z-score; one-way ANOVA: $F(3,8)=1.008$, $p=0.4381$] or timing [delay time of centre of mass; one-way ANOVA: $F(3,8)=1.186$, $p=0.3745$]. n=4 (WT), 3 (*App*$^{NL\text{-}G\text{-}F}$, Oligo-KO), 2 (Neuron-KO) mice.

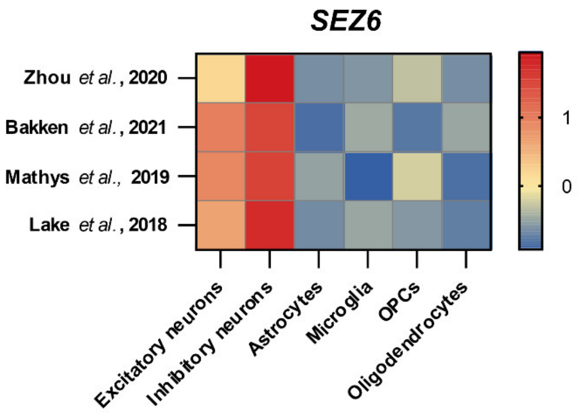


**Supplementary Fig. 15: Seizure 6 protein (*SEZ6*) is expressed at high levels in neurons, but not in oligodendrocytes.** Heatmap showing the $\log_2$ (norm count) z-score of *SEZ6* across different cell types from 3 different datasets[7–10]. [OPCs: oligodendrocyte precursor cells].
